# Host-generalist fungal pathogens of seedlings may maintain forest diversity via host-specific impacts and differential susceptibility among tree species

**DOI:** 10.1101/2020.12.24.424362

**Authors:** Erin R. Spear, Kirk Broders

**Affiliations:** Smithsonian Tropical Research Institute, Panama City, Panama, Republic of Panama

**Keywords:** *Clonostachys*, host generalists, host-specific impacts, multi-host pathogens, *Mycoleptodiscus*, Nectriaceae, seedlings, tropical forests

## Abstract

- Host-specialized pathogens are credited with the maintenance of tropical forest diversity under the Janzen-Connell hypothesis. Yet, in diverse forests, selection may favor pathogens with broad host ranges given their passive dispersal and the relative rarity of tree species.
- We surveyed the host associations of potential pathogens isolated from symptomatic seedlings in the forests in Panama and used inoculations to assess: (i) the pathogenicity and host ranges of 27 fungal isolates, and (ii) differences among tree species in susceptibility.
- Thirty-one of the 33 non-singleton OTUs isolated from seedlings are multi-host. All 31 multi-host OTUs exhibit low to moderate specialization, and we observed phylogenetically overdispersed host use for 19 OTUs. The pathogenicity of 10 isolates was experimentally confirmed; nine caused disease in seedlings in multiple families. However, the outcome of infection differs among tree species susceptible to a given multi-host pathogen. Furthermore, some tree species were seemingly resistant to all fungi tested, while others were susceptible to multiple fungi. Tree species adapted to environments with lower disease pressure were most likely to exhibit disease.
- Our results suggest that generalist pathogens contribute to the maintenance of local and regional forest diversity via host-specific impacts and the exclusion of disease-sensitive trees from disease-prone habitats.

## Introduction

The maintenance of local diversity is commonly attributed to host-specific natural enemies believed to prevent any single host species from becoming competitively dominant. Under the Janzen-Connell hypothesis (JCH), conspecific seeds and seedlings at high densities near conspecific adults suffer disproportionately higher mortality relative to heterospecific seeds and seedlings due to an accumulation of host-specialized enemies (Gillett, 1962; Janzen, 1970; Connell, 1971; Comita *et al*., 2014). However, support for the JCH has often been based on spatial and temporal patterns of conspecific negative density-dependent mortality (e.g., Harms *et al*., 2000; Comita *et al*., 2010), which cannot identify the mechanism(s) driving the observed patterns.

Experimental studies suggest that pathogens may contribute to density- and distance-dependent mortality of conspecific seedlings (Augspurger & Kelly, 1984; Bell *et al*., 2006; Mangan *et al*., 2010; Bagchi *et al*., 2014). However, the pathogens are rarely identified, and most studies fail to experimentally address a crucial assumption of the JCH, that the pathogens exhibit high host specificity with regard to locally available hosts (but see Packer & Clay, 2000; Augspurger & Wilkinson, 2007; Liu *et al*., 2012).

Furthermore, the relative rarity of tree species in diverse tropical forests and the passive dispersal of fungi may select for pathogens with intermediate to broad host ranges (May, 1991; Coley & Barone, 1996). Indeed, mounting evidence from tropical and temperate forests suggests that multi-host pathogens are prevalent (Augspurger & Wilkinson, 2007; Gallery *et al*., 2007; Gilbert & Webb, 2007; Kluger *et al*., 2008; Hersh *et al*., 2012; Schweizer *et al*., 2013; Sarmiento *et al*., 2017; Chen *et al*., 2019).

Pathogens are classified as specialists or generalists based on the number and phylogenetic relatedness of host species (Barrett *et al*., 2009). Host-specialized multi-host pathogens are predicted to exhibit a phylogenetic signal to their host range (Barrett *et al*., 2009). A variety of plant traits influence the outcome of plant-pathogen interactions, specifically susceptibility to infection, and resistance and tolerance post-infection (Coley & Barone, 1996; Bradley *et al*., 2003; Barrett *et al*., 2009). Plant defense traits are often, but not always, phylogenetically conserved (Fine *et al*., 2006; Agrawal, 2007; Kursar *et al*., 2009). Assuming that closely related plants are more likely to possess similar defense traits, the likelihood of tree species sharing a pathogen should decrease with decreasing phylogenetic relatedness (Barrett *et al*., 2009). Pathogens specialized at the clade-level could still maintain plant community diversity, but at higher taxonomic levels (Parker *et al*., 2015).

While specialist pathogens are invoked under JCH, generalist pathogens may contribute to the maintenance of host community diversity via effective specialization (Benítez *et al*., 2013). One form is host-specific impacts (Sarmiento *et al*., 2017), meaning disease severity differs among hosts of a given pathogen. Host-specific impacts could be an equalizing mechanism that maintains plant community diversity if the growth and survival of infected seedlings varies among host species (Barrett *et al*., 2009); particularly, if differential impacts reflect ecological trade-offs between allocation of resources to defense, competitive ability, and/or tolerance of stressors. For example, for tree species adapted to environments with high and/or consistent disease pressure, the fitness benefits of physical and chemical resistance and/or tolerance traits may outweigh the associated allocation costs of those traits, while the costs may outweigh the benefits for tree species adapted to environments with low and/or inconsistent disease pressure (Haak *et al*., 2012). The risk of pathogen attack, referred to here as pathogen pressure, varies with environmental conditions that impact pathogen and plant fitness (Barrett *et al*., 2009). High pathogen pressure has been observed in areas with low light, high soil moisture, and persistent fog and dew (Augspurger & Kelly, 1984; Bradley *et al*., 2003; Mordecai, 2012; Spear *et al*., 2015). Plants adapted to environments with high and/or consistent pathogen pressure may be under selection for increased defenses, whereas plants adapted to low and/or inconsistent pathogen pressure environments may be poorly defended (Coley & Barone, 1996, Talley *et al*., 2002).

Here we describe (i) pathogens infecting tree seedlings in the forests of Panama, (ii) their host ranges, (iii) interspecific variability among tree species in pathogen susceptibility, and (iv) the relationship between disease susceptibility and plant habitat associations. We had four central hypotheses. First, in these diverse tropical forests, seedling pathogens are often multi-host. Second, there is a phylogenetic signal to the host ranges of these multi-host pathogens (i.e., pathogen sharing is positively correlated with host relatedness). Third, tree species differ in their susceptibility to pathogens because the cost-to-benefit ratio of defense traits differs among species (Strauss & Agrawal, 1999; Endara & Coley, 2010). Fourth, plants adapted to environments with low and/or inconsistent disease pressure are more susceptible to disease than those adapted to high and/or consistent disease pressure.

## Materials and Methods

### Identification of putative pathogens

Potentially pathogenic fungi and oomycetes were isolated from 124 seedlings, representing 26 tree species in 13 families, with disease on the leaf, stem, and/or root (Supporting Information Fig. **S1;** Table **S1**). Symptomatic seedlings were collected from seven forest sites in 2007, 2010-2012, and 2019 by: (1) opportunistic collection of naturally occurring seedlings (*n* = 59); (2) collection of seedlings germinated in a shadehouse and then transplanted to forest sites (*n* = 22), and (3) collection of seedlings that germinated from surface-sterilized seeds planted directly in the forests (*n* = 43) (Table **S2**). Most putative pathogens were isolated from leaves (52.13%), followed by stems (37.44%) and roots (10.43%). Molecular identification was completed by bidirectional Sanger sequencing of either the nuclear ribosomal Internal Transcribed Spacer (ITS) region (149 isolates) or the ITS plus an adjacent portion of the large subunit (ITS+LSU) (62 isolates). See Table **S2** and Spear (2017) for details.

All 211 unique isolates were identified and assigned to operational taxonomic units (OTUs) based on 99% sequence similarity (minimum overlap of 40 bp) in Sequencher v. 5.4.6. We used a stringent similarity threshold that splits rather than lumps, while accounting for ≤1% sequencing error, because we make statements about host specificity and to avoid masking strain-specific effects (Higgins *et al*., 2011). We estimated the taxonomic placement of the fungal isolates by querying all 209 edited sequences against the well-curated Full UNITE+INSD dataset for Fungi (v. 8.2, released 2020-02-04; Abarenkov *et al*., 2020) (detailed in Methods **S1**). For the two oomycetes, we queried their sequences against GenBank (accessed 8-Oct-2020; Benson *et al*., 2012) and the curated database *Phytophthora*-ID (v. 2.0, accessed 8-Oct-2020; Grünwald *et al*., 2011). All assigned nomenclatures represent estimates as: (i) the ITS region is an imprecise barcode for species discrimination for certain taxa; (ii) multi-locus phylogenies are required for accuracy; and (iii) precision is limited by incomplete taxonomic and geographic coverage of databases (Kang *et al*., 2010; Hofstetter *et al*., 2019; Lücking *et al*., 2020). Additionally, sequences in the UNITE database are clustered into hypothesized species-level groups (Species Hypotheses, SHs) based on sequence similarity thresholds ranging from 97–100% (Kõljalg *et al*., 2013; Robbertse *et al*., 2017), while we used 99% sequence similarity to designate OTUs because 97-98.5% is too relaxed for species delimitation for some taxa (Lücking *et al*., 2020). Because our grouping strategy often differed from that of the UNITE database, multiple, distinct OTUs in our study sometimes share the same nomenclature based on the SH of the best-matching UNITE reference sequence (e.g., OTUs Z and AAG are both annotated as *Colletotrichum brevisporum*). Finally, for certain groups of fungi, UNITE SHs erroneously include distinct species (Robbertse *et al*., 2017).

Sequences for the 90 previously published fungal isolates (Spear, 2017) were submitted to the NCBI GenBank database (KY413686–KY413775). Sequences for the 121 isolates collected in 2019 will be submitted to GenBank (XXXXXXX– XXXXXXX).

To characterize sampling efficacy and the relative abundance of OTUs, we calculated the richness of observed OTUs and plotted the taxon accumulation curve (vegan package; Oksanen *et al*., 2019) and rank-abundance distribution (BiodiversityR package; Kindt & Coe, 2005) (*n* = 211 isolates from 124 seedlings). We defined OTUs observed ≥10 times as abundant.

### Survey-based assessments of host associations and ranges of putative pathogens

Our survey-based assessments of the host associations and ranges of putative pathogens utilize our full dataset (*n* = 211 isolates, representing 66 OTUs, from 26 tree species) and two subsets to ameliorate the influence of undersampled taxa. Subset A excludes singleton OTUs (*n* = 178 isolates, representing 33 OTUs, from 24 tree species) and B excludes singleton and doubleton OTUs and tree species (*n* = 144 isolates, representing 22 OTUs, from 13 tree species). All three datasets include isolates collected in all 5 yr.

We used four approaches to characterize host associations. We (1) summed and plotted the number of observed host tree species (unweighted links) (full dataset), and (2) calculated Blüthgen’s weighted specialization index d’ for each of the 33 non-singleton OTUs (subset A; bipartite package; Blüthgen *et al*., 2006; Dormann *et al*., 2008). The d′ index is useful for identifying discriminatory host use because it accounts for interaction strength (i.e., strong, frequent vs. weak, occasional interactions) and sampling intensity, while the number of interacting partners does neither (Blüthgen *et al*., 2006; Dormann, 2011). The index estimates specialization by calculating the degree to which observed differs from expected host use given partner availability (Blüthgen *et al*., 2006; Dormann, 2011). We could not adjust for the relative abundances of tree species when calculating d′ as the tree species are not uniformly distributed across the seven forest sites (Pyke *et al*., 2001). Thus, we relied on the interaction matrix’s marginal sums (column- and row-wise totals) as a measure of partner availability, meaning that, for our data, a generalized microbe infects tree species proportional to their sampling intensities, while a specialized microbe infects tree species independent of their sampling intensities (Blüthgen *et al*., 2006). We considered d’ values ranging from 0–0.33, 0.34–0.67, and 0.68–1.00 to represent low, moderate, and high specialization, respectively (Blüthgen *et al*., 2006; Spear & Mordecai, 2018). We (3) calculated the observed number of shared OTUs (rich package; Rossi, 2011) and, because that metric generally results in an underestimation of similarity due to undetected species, we (4) estimated similarities in the communities of putative pathogens shared between pairs of tree species via the inverse of the Chao dissimilarity index (vegan package; Oksanen *et al*., 2019) (full dataset and subset B). The Chao index accounts for the relative abundances of observed OTUs and unseen shared OTUs, making it relatively robust to unequal sampling and diverse guilds with many rare taxa (Chao *et al*., 2005).

To determine if there is a phylogenetic signal to the host ranges of the OTUs, (i) we used mean pairwise phylogenetic distance (MPD) to determine if a given multi-host OTU infects phylogenetically clustered or dispersed hosts, and (ii) we used two binomial generalized linear models (GLMs) (logit link) to determine whether putative pathogen community similarity is positively correlated with host relatedness. Specifically, for the 31 OTUs isolated from ≥2 tree species, we compared the observed MPD between tree species infected by a given OTU to the MPD expected under a null model with random host-OTU associations (full dataset; 999 randomizations of the host phylogeny tip labels; picante and ape packages; Kembel *et al*., 2010; Paradis & Schliep, 2018). For the GLMs, we used our full dataset and subset B, and we regressed and plotted the pairwise estimated community similarities (1 - Chao index) against the pairwise phylogenetic distances. The phylogeny of tree species was generated by Phylomatic (phylodiversity.net/phylomatic/), which called the Zanne *et al*. (2014) supertree.

We tested for correlations between (i) the frequency with which a given fungal OTU was isolated and the number of tree species from which it was isolated, and (ii) sampling intensity (number of seedlings collected) and the number of OTUs isolated from a given tree species using non-parametric Spearman’s rank tests (full dataset; stats package; R Core Team, 2020). One-tailed tests were used because we had directional, literature-based *a priori* hypotheses (Quinn & Keough, 2002; Ruxton & Neuhäuser, 2010) that: (i) the most abundant fungal taxa would be multi-host, and (ii) observed fungal richness associated with a given plant species would increase with increased sampling (Ferrer & Gilbert, 2003; Spear & Mordecai, 2018).

### Experimental assessments of pathogenicity and host range

In 2011 and 2012, inoculation experiments were used to assess the pathogenicity and host specificities of 27 fungal isolates collected in 2010 and 2011 (Fig. **S2a**). We tested the isolates, representing 18 OTUs, against seedlings of 35 tree species, representing 20 families (Table **S1**). We could not test four isolates against the tree species from which they were originally isolated because seeds were unavailable or available in insufficient quantities (e.g., ERS033; Fig. **3**; Table **S1**). Related, recreating the exact conditions under which disease developed for a specific fungus-plant combination is challenging. Thus, some tree species were included as phytometers of pathogenicity based on previously observed disease susceptibility (e.g., *Luehea seemannii*, Augspurger & Wilkinson 2007; *Dalbergia retusa*, Spear, personal observation). Tree species-by-isolate combinations were not fully reciprocal (285 of 945 possible combinations; 3,154 seedlings total) due to space, time, and seed availability constraints. A given fungal isolate was tested against 3–28 tree species and 48% of isolates evaluated were tested against seedlings of ≥10 tree species. Because an objective of the experiments was to describe pathogen host range, once isolates displayed pathogenicity, we prioritized testing those isolates against additional tree species (12-28 tree species, median 15) over expanding the testing of isolates failing to generate significant disease. A given tree species was tested against 1–26 isolates, and 29% of tree species were tested against ≥10 isolates.

Our inoculation technique simulated infested plant material in the soil to passively inoculate the seedlings with the fungal isolates selected for screening (modified from Augspurger & Wilkinson, 2007). To generate seedlings for the experiments, seeds were collected from forests bordering the Panama Canal (May-Nov. 2011, May-June 2012), surface-sterilized (sequential washes of 95% EtOH [10 s], 10% commercial bleach [2 min], 70% EtOH [2 min]), and germinated in flats of autoclave-sterilized (1 hour at 121° C, twice) commercial soil (Fig. **S2b**). Seedlings were transplanted to individual pots containing autoclaved commercial soil and either rice visibly colonized by one of the isolates or inoculum-free, autoclave-sterilized rice (control/sham treatment) (Fig. **S2c-f**). Depending on seed availability, 5–12 seedlings per species were included in each treatment.

Inoculation experiments were conducted in Smithsonian Tropical Research Institute shadehouses in Gamboa, Panama during the wet seasons of 2011 and 2012 (Fig. **S2a**). The two screened-in shadehouses were covered on all sides with shadecloth and their roofs were covered with plastic to minimize external contamination and to mimic the forest understory. Shadehouse air temperatures (26.1 and 25.6°C) did not significantly differ from those recorded simultaneously in a nearby forest understory (25.4 and 25.3°C; Paton, 2019a). Relative humidity was significantly lower in the shadehouses (85.1 and 87.6%) than in the forest understory (95.6 and 96.7%; Paton, 2019b). Photosynthetically active radiation (PAR) reaching the shadehouse seedlings was slightly higher (1.4 and 1.7% of full PAR) than observed in a nearby forest understory (1.3%; Brenes-Arguedas *et al*., 2011). See Table **S3** for details. Light was somewhat variable within a given shadehouse. To counteract this potentially confounding factor, seedlings were rotated among the tables weekly.

Disease was documented every 3 d (for ≤5 wk) and categorized as: (i) mortality; (ii) stem damage (Fig. **S2g-o**); (iii) wilting (Fig. **S2p**,**q**); and (iv) stunting (Fig. **S2r**,**s**). A seedling was classified as morbid if it had any of the four symptoms. Wilting was considered a disease symptom because the soil did not dry out (seedlings were hand-watered every 3 d) and pathogens attacking roots can impede water uptake (Yadeta & Thomma, 2013). Similarly, stunting can indicate pathogen-caused damage to roots and/or the vascular system (Yadeta & Thomma, 2013). Additionally, stunting may occur because energy and resources are allocated to defenses and/or tissue replacement rather than growth (Huot *et al*., 2014). We did not document root damage and could not ascribe foliar damage to a specific treatment because we observed a synchronous occurrence of morphologically similar foliar necrosis across distinct treatments and the controls, suggesting that the causative pathogen (*Xylaria* sp., unpublished data) blew into the shadehouses. Thus, neither damage type was included in our analyses.

To compare the proportion of symptomatic seedlings in the inoculated versus inoculum-free control treatments, we used generalized linear models (GLMs) with binomial error distributions (probit link) and mean bias-reducing score adjustments (brglm package; Firth, 1993; Kosmidis & Firth, 2009; Kosmidis, 2019). We used bias-reduced GLMs because we observed complete and quasi-complete separation of data for some isolate-by-tree species combinations (i.e., treatment perfectly, or near perfectly, predicted the binary outcome; Albert & Anderson, 1984). The response variables were presence/absence of (i) general morbidity and (ii) mortality. Each isolate-by-tree species combination was considered individually for each response variable. We considered an isolate pathogenic if, for at least one tree species, a significantly greater proportion of inoculated seedlings were symptomatic than paired control seedlings (using α = 0.10 because small sample sizes limited our power to detect treatment effects; Quinn & Keough, 2002; Thiese *et al*., 2016). We tested for a correlation between the frequency with which a fungal OTU was isolated and the number of tree species in which it caused statistically significant disease using a non-parametric Spearman’s rank test. We used a one-tailed test because we had the *a priori* hypothesis that the most abundant fungal taxa would be multi-host.

### Plant traits and disease susceptibility

We used logit-link beta-binomial generalized linear models to explore disease susceptibility as a function of habitat associations and seed size (aod package; Lesnoff & Lancelot, 2012). We obtained data for seed size, shade tolerance, and spatial distribution relative to annual rainfall from published sources (Table **S1**). In all models, the response variable was presence/absence of morbidity in inoculated seedlings, controlling for the morbidity observed in paired control seedlings and considering all 27 isolates tested (pathogens and non-pathogens alike). Because negative relative proportions are biologically equivalent to zero (no disease caused), and for statistical purposes, we converted all negative values to zero. We used two global models that contained (i) the additive main effects of seed size (continuous) and shade tolerance (categorical), and their interaction, or (ii) the additive main effects of seed size and distribution relative to rainfall (categorical), and their interaction. We excluded seeds with average dry masses >3,000 mg to improve model stability. For both categorical predictors, we limited our comparisons to binary extremes (i.e., only shade-intolerant and -tolerant, and wet- and dry-site tree species). We analyzed disease as a function of shade tolerance (26 tree species, 179 observations: 42 for intolerant, 137 for tolerant spp.) and spatial distribution (15 tree species, 154 observations: 108 for dry-site, 46 for wet-site spp.) separately due to differences in the availability of trait data. For the two sets of models, automated model selection and averaging was done based on the global models, the Akaike information criterion with a correction for small sample sizes (AICc), and ΔAICc ≤ 2 (MuMIn package; Bartoń, 2020).

All data were processed in R v. 4.0.0 (R Core Team, 2020). In addition to the aforementioned packages, we used fossil (Vavrek, 2011), ggplot2 (Wickham, 2016), otuSummary (Yang, 2020), plot.matrix (Klinke, 2019), plyr (Wickham, 2011), reshape2 (Wickham, 2007), and VennDiagram (Chen, 2018) for data formatting and figures.

## Results

### Abundances and taxonomic assignments of putative pathogens

The 211 isolates represent 66 operational taxonomic units (OTUs); however, the taxon accumulation curve is non-asymptotic (Fig. **S3b**). Most of the OTUs are relatively rare, 33 are singletons (Figs **1**, **S3a**; Table **S4**). Four OTUs are common (observed ≥10) and comprised 35% of all isolates: A–*Colletotrichum xanthorrhoeae*, B–Nectriaceae, C– *Mycoleptodiscus suttonii*, and F–*Diaporthe tulliensis* (Figs **1**, **S3a**; Table **S4**). The 209 fungal and two oomycete isolates represent five classes (Sordariomycetes, Dothideomycetes, Agaricomycetes, Eurotiomycetes, Oomycetes) and 11 orders. Most (81.5%) of the isolates could be assigned to 26 genera; *Colletotrichum, Diaporthe*, and *Mycoleptodiscus* were most frequently isolated (Fig. **1**; Table **S4**). We observed overlap in fungal OTUs among the tissues sampled: leaves, stems, and roots (Fig. **S4d**). While sampling occurred over 5 yr, there was overlap in the tree species and forest sites sampled, and the isolation and molecular techniques used (Table **S2**). Accordingly, we observed overlap in fungal OTUs among the years, seedling collection methods, and isolation media (Fig. **S4a-c**).

**Figure 1.**
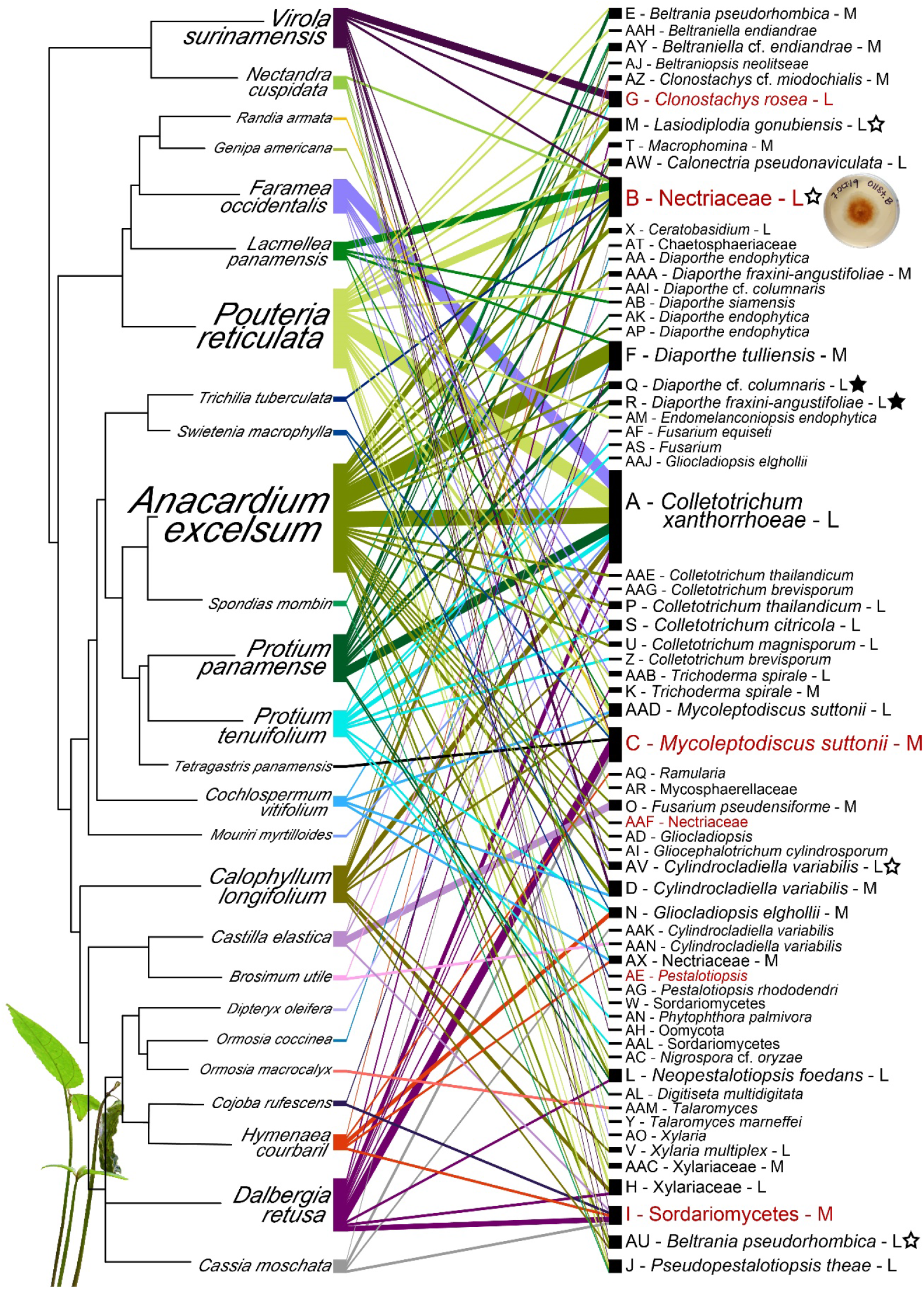
Host associations of the fungi and oomycetes isolated from symptomatic seedlings (full dataset: *n* = 211 isolates). The 26 host tree species (left) are arranged according to their phylogenetic relationships. The estimated taxonomic placement of each of the 66 microbial OTUs (right) is preceded by a unique letter identifier. Because our OTU grouping strategy (99% sequence similarity) often differed from that of the UNITE database, distinct OTUs in our study sometimes share the same nomenclature based on the species hypothesis of the best-matching reference sequence in UNITE (e.g., Z and AAG). For the non-singleton OTUs, nomenclature is followed by an L or M, indicating low or moderate estimated specialization (d’). The width of each line represents the number of times a given OTU was isolated from seedlings of a given tree species. For both tree species and OTUs, font size is scaled to the number of interactions. OTUs with significant and marginally significant phylogenetic patterns of host use (Table S4) are marked with stars, solid indicating clustered and open indicating overdispersed. The six fungal OTUs with members that exhibited statistically significant (α = 0.10) pathogenicity during the inoculation experiments are listed in red.

### Fungi associated with symptomatic seedlings are often host generalists

The non-singleton fungi isolated from symptomatic seedlings are predominately multi-host generalists with wide host ranges. Of the 33 non-singleton OTUs, 31 were isolated from 2–9 tree species (median = 3) (Fig. **1**; Table **S4**). The number of tree species from which a fungal OTU was isolated is positively correlated with isolation frequency (Spearman’s rank correlation: rho = 0.96, *P* < 0.001; blue points Fig. **S5**). A–*C. xanthorrhoeae* was isolated with the greatest frequency (35 times) and from the greatest number of tree species (nine in seven families) (Fig. **1**; Table **S4**). Only two of the non-singleton OTUs (T–*Macrophomina*, X–*Ceratobasidium*) were not multi-host, but each was isolated only twice, and from the best-sampled tree species, suggesting their observed host ranges are a sampling artifact (Fig. **1**; Tables **S1, S4**). Based on the specialization index d’, 57.6% of 33 the non-singleton OTUs, including A–*C. xanthorrhoeae* and B–Nectriaceae, exhibit low host-specialization and 42.4%, including C–*M. suttonii* and F–*D. tulliensis*, exhibit moderate host-specialization at the species level (Table **S4**). We observed phylogenetically overdispersed host use for 19 of the 31 multi-host OTUs, significant for B–Nectriaceae and AV–*Cylindrocladiella variabilis*, and marginally significant for M–*Lasiodiplodia gonubiensis* and AU–*Beltrania pseudorhombica* (Fig. **1**; Table **S4**). Conversely, 12 OTUs exhibited phylogenetically clustered host use, marginally significant for R–*Diaporthe fraxini-angustifoliae* and Q– *Diaporthe* cf. *columnaris* (Fig. **1**; Table S**4**). Both OTUs were solely isolated from *Anacardium excelsum* (Anacardiaceae, Sapindales) and *Protium panamense* (Burseraceae, Sapindales), providing tentative evidence of a phylogenetic signal to host range.

### Tree species are infected by various microbes, and their microbial communities overlap

Multiple OTUs were isolated for all tree species with >2 seedlings collected (Fig. **1**; Table **S1**). Predictably, the number of OTUs isolated from a given tree species is strongly positively correlated with sampling intensity (Spearman’s rank correlation: rho = 0.95, *P* < 0.001; Fig. **1**). The greatest richness of fungi (24 OTUs) was recovered from the best-sampled tree species (41 isolates), *A. excelsum* (Table **S1**).

We observed overlap among the microbial communities of distantly related tree species (Fig. **1**). Based on subset B, only tree species and OTUs observed ≥3 times, the estimated average similarity (1 - Chao index) between the microbial communities of pairs of tree species is 0.21 (median = 0.15; Table **S5**). The GLMs suggest microbial community similarity is unrelated to phylogenetic distance between tree species (Fig. **2**; full dataset: Est. ± SE = 0.002 ± 0.003, *z*-value = 0.60, *P* = 0.55; subset B: Est. ± SE = 0.002 ± 0.004, *z*-value = 0.36, *P* = 0.72).

**Figure 2.**
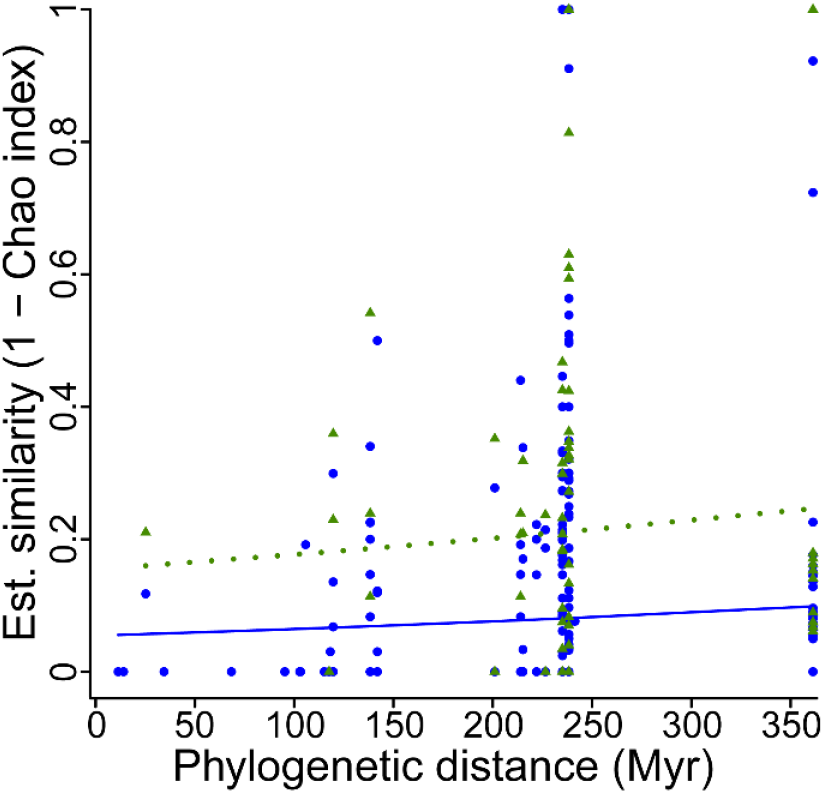
Estimated pairwise similarities of OTU communities are unrelated to pairwise phylogenetic distances (time of independent evolution in Myr). The blue circles and solid line (GLM: Est. ± SE = 0.002 ± 0.003, *z*-value = 0.60, *P* = 0.55) are based on the full dataset, while the green triangles and dotted line (GLM: Est. ± SE = 0.002 ± 0.004, *z*-value = 0.36, *P* = 0.72) are based on subset B, only tree species and OTUs observed ≥3 times.

### Identities and impacts of confirmed pathogens

We experimentally confirmed the pathogenicity of 10 of the 27 fungal isolates tested (Fig. **3**). The pathogenic isolates belong to six OTUs, all in the phylum Ascomycota, and were isolated from stems, roots, and leaves of seedlings of seven tree species in seven families (Fig. **3**; Table **1**). Based on morbidity (i.e., any of the four symptoms: wilting, stunting, stem damage, or mortality), significant pathogenicity was observed for 20 of the 285 tree species-by-isolate combinations (Fig. **3**; Table **1**). For 13 of these combinations, infection led to mortality (Table **1**). Multiple symptoms were often observed for a single tree species-by-pathogen combination (e.g., *Annona glabra* seedlings suffered stem damage, wilt, and mortality when inoculated with isolate ERS029). Seven of the pathogenic isolates were tested against the tree species from which they were originally isolated, but for those tree species-by-isolate combinations neither morbidity nor mortality significantly differed between the inoculated and control seedlings (Fig. **3**).

**Table 1.**
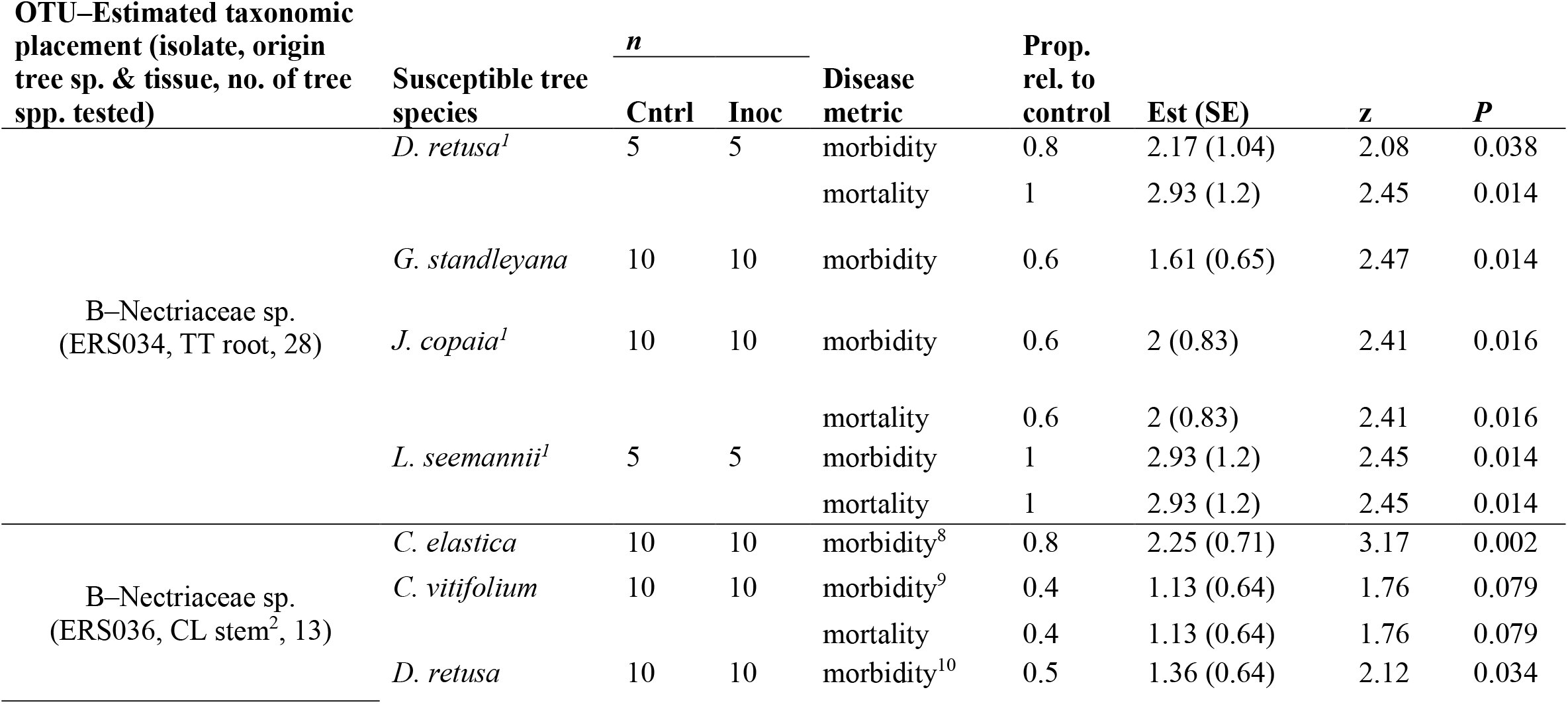
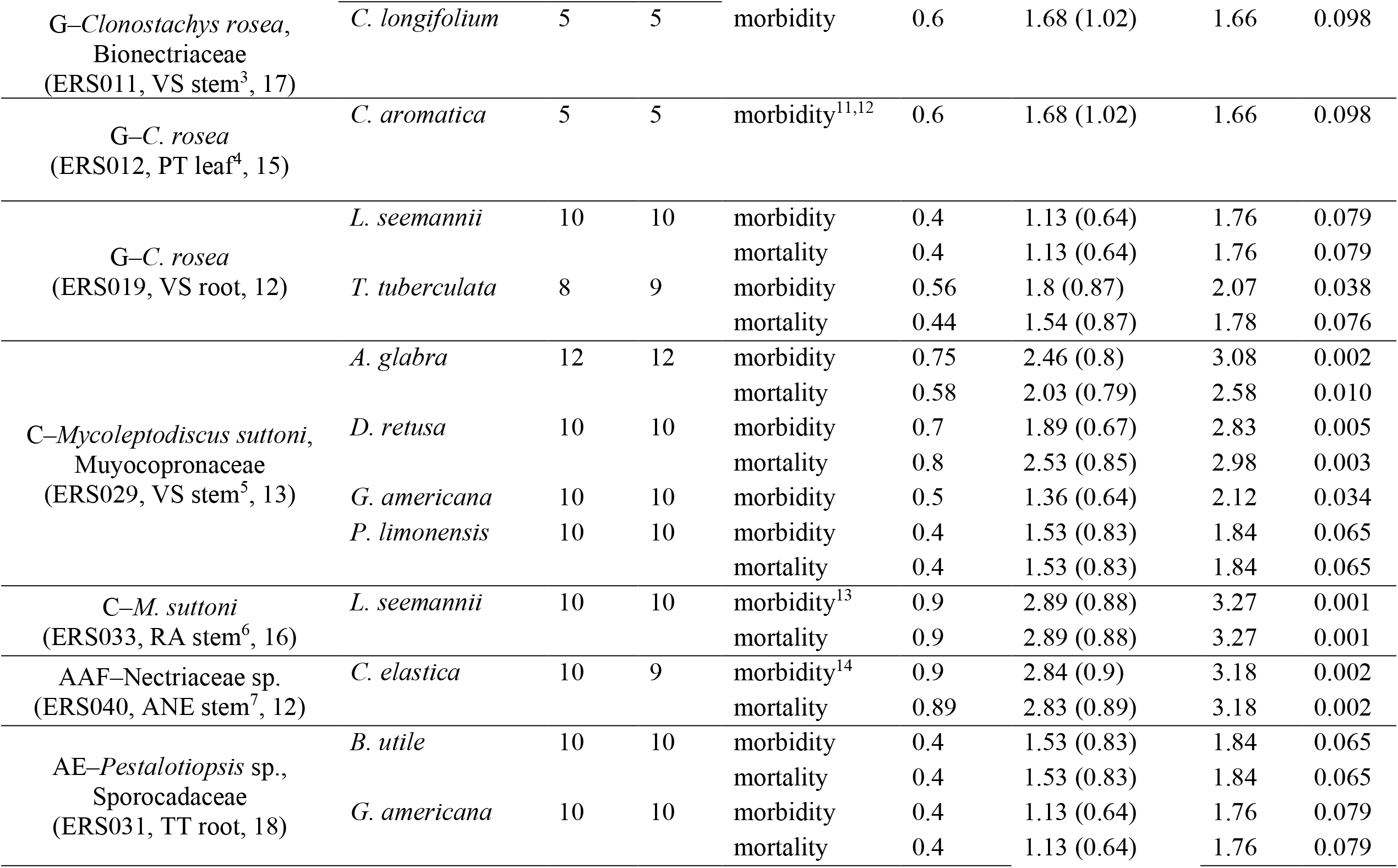
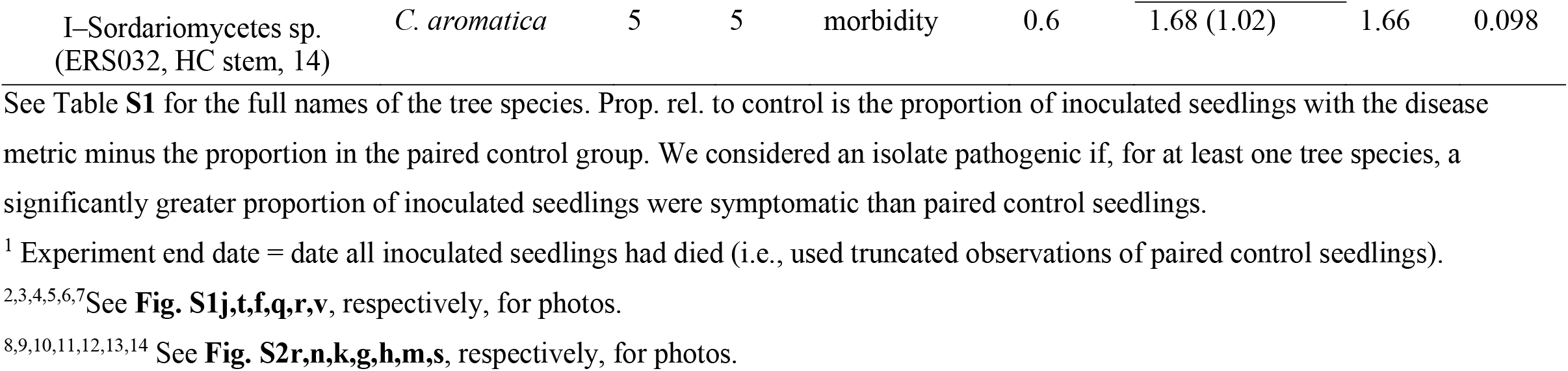
The 20 fungal isolate-by-tree species combinations for which significant pathogenicity was observed based on the presence/absence of (i) general morbidity and (ii) mortality in the inoculated versus control seedlings, analyzed with bias-reduced generalized linear models assuming binomial error distributions (probit link) and using α = 0.10 because small sample sizes limited our power to detect treatment effects.

**Figure 3.**
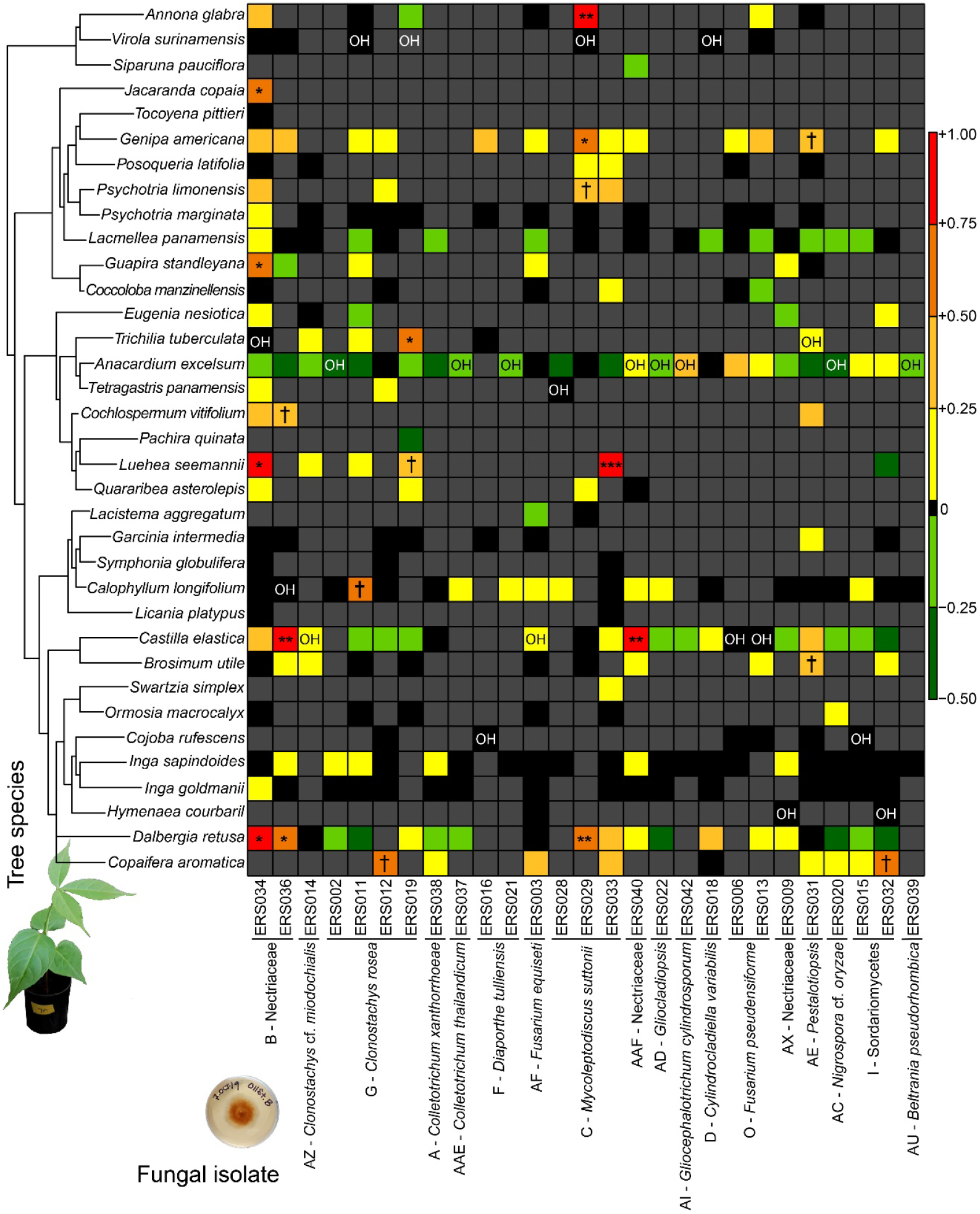
Pathogenicity and host ranges of the 27 fungal isolates tested, and susceptibility of the 35 tree species tested, based on seedling morbidity. Fungal isolates are grouped according to their OTU (horizontal axis). Tree species are arranged based on their phylogenetic relationships (vertical axis). OH (original host) denotes the tree species from which an isolate originated. Tree species-by-isolate combinations for which evidence of pathogenicity was observed (proportion of inoculated seedlings with symptoms > proportion in paired control group) are depicted in shades of yellow (pale for minimal) to red (dark for substantial). Combinations for which there was no evidence of pathogenicity (proportion of inoculated seedlings with symptoms ≤ proportion in paired control group) are depicted in black and shades of green. Gray cells represent an untested combination. For 20 of the 285 combinations tested, we observed significant pathogenicity (***, *P* ≤ 0.001; **, *P* ≤ 0.01; *, *P* ≤ 0.05; †, *P* ≤ 0.10; Table **1**).

### Fungi with confirmed pathogenicity are multi-host

We show that fungal pathogens can attack tree species belonging to phylogenetically dispersed families. Five of the 10 pathogenic isolates successfully attacked multiple tree species (Table **1**). The 10 pathogenic isolates generated significant disease in 6–31% of the tree species against which they were tested (median = 10%). At the high end, ERS029 (C–*Mycoleptodiscus suttonii*) caused significant disease in 31% of the 13 tree species it was tested against, and the four susceptible tree species belong to three orders (Fig. **3**; Tables **1, S1**).

Three of the OTUs were represented by multiple isolates: B–Nectriaceae, G– *Clonostachys rosea*, and C–*M. suttonii*. The two isolates of B–Nectriaceae were recovered from *Trichilia tuberculata* and *Calophyllum longifolium*, and combined were pathogenic on six of the 28 tree species tested (Fig. **3**; Table **1**). The three isolates of G– *C. rosea* were isolated from *Virola surinamensis* and *Protium tenuifolium*, and combined caused significant disease on four tree species (Fig. **3**; Table **1**). The two isolates of C–*M. suttonii* were isolated from *V. surinamensis* and *Randia armata*, and combined caused significant disease on five species (Fig. **3**; Table **1**). While the tree species-by-isolate combinations were not fully reciprocal, limiting conclusions, we observed possible strain-specific effects (e.g., B–Nectriaceae: ERS034 caused significant disease on *Guapira standleyana* while ERS036 did not; Fig. **3**). Finally, there is a marginally significant positive association between the frequency with which a fungal OTU was isolated and the number of tree species in which it caused statistically significant (α = 0.10) disease (Spearman’s rank correlation rho = 0.72, *P* = 0.053; green points Fig. **S5**).

### Tree species are differentially impacted by pathogens

We documented interspecific differences in disease susceptibility and impact among tree species. Most tree species tested were seemingly resistant to, or tolerant of, all isolates tested, suffering minimal to no fitness impacts despite soil inoculation (Fig. **3**). For 13 of the 35 tree species tested, at least one of the 27 fungal isolates tested caused significant disease (Table **1**). Five tree species experienced significant disease from 2–3 of the isolates tested (Table **1**). Those five species experienced significant disease from 10–50% (median = 15%) of the isolates against which they were tested (Fig. **3**).

The 13 susceptible tree species differed in their tolerance of attack. For most of the susceptible tree species, pathogen attack did not consistently lead to death, the impact from infection depended on the pathogenic isolate with which it was inoculated. For example, while *Castilla elastica* seedlings suffered significant disease when inoculated with two isolates (ERS036, ERS040), they only suffered significant mortality when inoculated with ERS040 (AAF–Nectriaceae) (Table **1**). Furthermore, for those tree species that were susceptible to the same multi-host pathogen, the impact from attack differed among species. For example, isolate ERS036 (B–Nectriaceae) caused significant morbidity on *C. elastica, Cochlospermum vitifolium*, and *Dalbergia retusa*, but only caused significant mortality on *C. vitifolium* (Table **1**).

### Plant traits and disease susceptibility

When considering all 27 fungal isolates tested (pathogens and non-pathogens), the proportion of morbid seedlings was influenced by a tree species’ seed size, spatial distribution relative to annual rainfall, and shade tolerance (Fig. **4a-d**; Tables **S6, S7**). Shade-intolerant tree species were more likely to exhibit disease than shade-tolerant species (*P* = 0.003; Fig. **4a,b**; Table **S6**). For both groups, the probability of disease decreases with increasing seed size, but to a greater extent for shade-intolerant species (*P* = 0.014; Fig. **4a,b**; Table **S6**). Tree species restricted to, or more abundant in, drier sites (dry-site) were more likely to exhibit disease than those restricted to, or more abundant in, wetter sites (wet-site) (*P* = 0.042; Fig. **4c,d**; Table **S7**). While we found no correlation between seed size and disease risk for wet-site species, disease risk decreases with increasing seed size for dry-site species (Fig. **4c,d**; Table **S7**).

**Figure 4.**
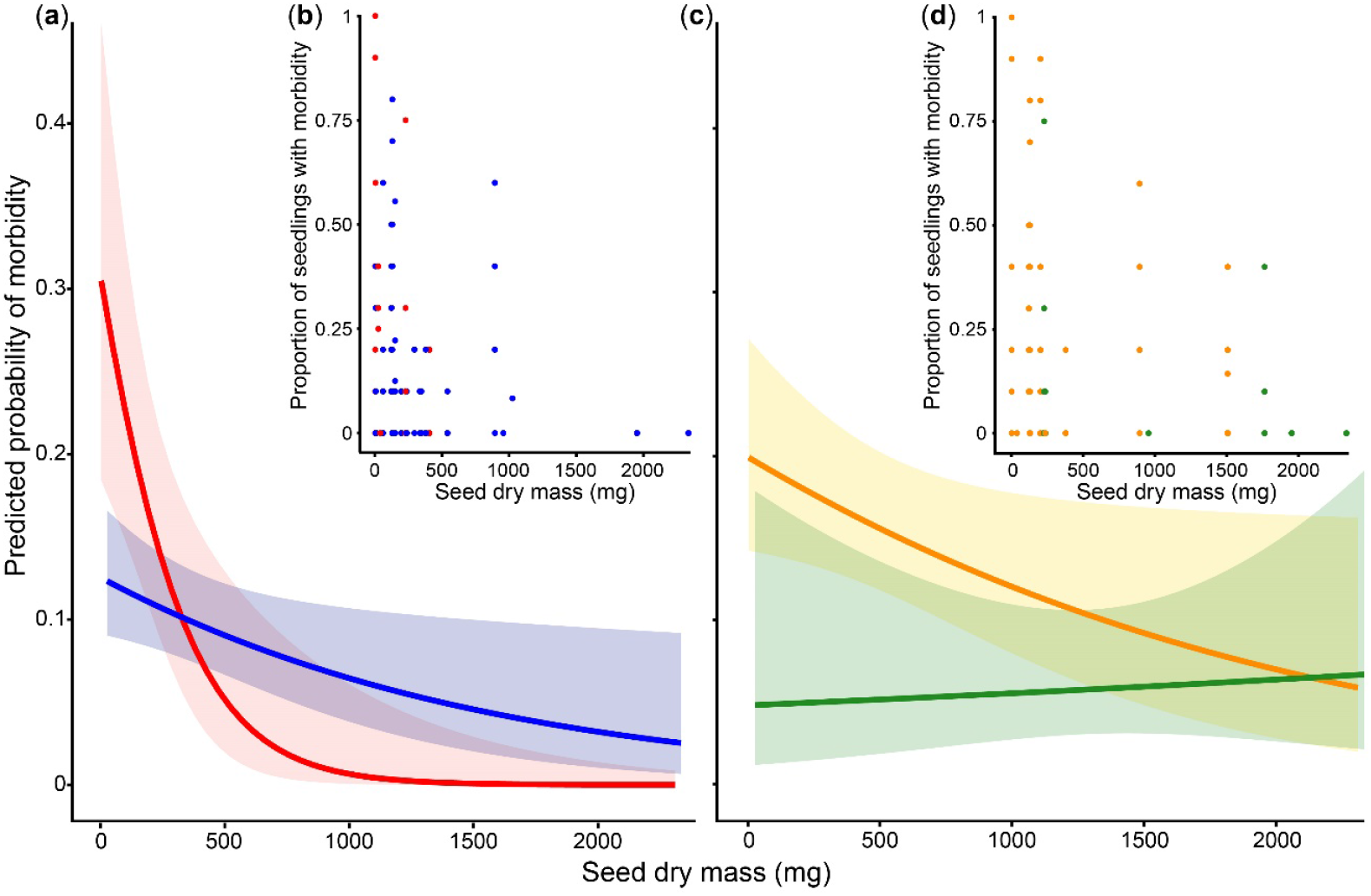
Relationships between disease susceptibility and seed size and shade tolerance (a,b), and seed size and spatial distribution relative to annual rainfall (c,d). Shade-intolerant tree species (red) are more likely to experience disease than shade-tolerant species (blue) (*P* = 0.003), and for both groups the probability of disease decreases with increasing seed size (a,b) (Table **S6**). Dry-site tree species (orange) are more likely to experience disease than wet-site species (green) (*P* = 0.042; Table **S7**). For dry-site species (orange) the probability of disease decreases with increasing seed size (c,d) (Table **S7**). The shaded areas represent 95% confidence intervals.

## Discussion

Pathogens are believed to shape the distributions and abundances of their hosts, effects often ascribed to host-specialized pathogens (Comita *et al*., 2014). However, there is increasing evidence that multi-host pathogens are prevalent in natural plant communities (Augspurger & Wilkinson, 2007; Gallery *et al*., 2007; Gilbert & Webb, 2007; Kluger *et al*., 2008; Hersh *et al*., 2012; Schweizer *et al*., 2013; Sarmiento *et al*., 2017; Spear & Mordecai, 2018). These multi-host pathogens may contribute to the maintenance of these communities via host-specific impacts (disease severity differs among a pathogen’s hosts), a form of effective specialization (Augspurger & Wilkinson 2007; Benítez *et al*., 2013; Sarmiento *et al*., 2017). Using a multifaceted approach, we identified fungal pathogens attacking seedlings in the forests of Panama, and show that they can attack phylogenetically dispersed hosts. Furthermore, we demonstrate that tree species differ in their susceptibility to pathogens, with trees adapted to environments with low and/or inconsistent disease pressure being more susceptible to disease. Collectively, these results suggest that generalist pathogens contribute to the maintenance of local and regional plant diversity.

### Pathogens and their impacts

We observed 66 OTUs, but only four were observed >10 times (Figs **1**, **S3a**; Table **S4**). The relative rarity of the OTUs, coupled with a non-asymptomatic taxon accumulation curve, indicate incomplete sampling and a diverse community of seedling-associated fungi (Fig. **S3**). The most frequently isolated fungal genera (e.g., *Colletotrichum*; Fig. **1**) and those with experimentally confirmed pathogenicity (e.g., *Clonostachys*; Table **1**) include previously identified phytopathogens (Nguyen *et al*., 2016).

Seven of the 10 pathogenic isolates caused significant mortality for at least one tree species during the inoculation experiments (Table **1**), reinforcing the importance of pathogens as a selective pressure and source of nonrandom seedling mortality that can directly shape plant community composition (Green *et al*., 2014). We also observed pathogen-caused stem damage, stunting, and wilting. These nonlethal infections can also be a structuring force by compromising a seedling’s ability to survive future stressors (Ditommaso & Watson, 1995).

Seventeen of the 27 fungal isolates tested appear to be nonpathogenic; however, some categorized as nonpathogenic may be pathogenic under optimal conditions. These isolates may not have caused disease in our experiments because we failed to: (1) test them against a susceptible host (each isolate was only tested against 3–28 tree species, a fraction of the species locally present); (2) test them against susceptible individuals of a host species, attributable to intraspecific genotypic variation in resistance; or (3) replicate the environmental conditions necessary for disease development (Barrett *et al*., 2009; Álvarez-Loayza *et al*., 2011). Additionally, while many of the fungi tested infect multiple tissue types (Table **1**; Fig. **S4d**), (4) some isolates deemed nonpathogenic based on root inoculation may only colonize leaves or stems. Experimental challenges two through four may also explain why the 10 pathogenic isolates did not generate significant disease on the host species from which they were originally isolated (Fig. **3**).

### Pathogens can infect phylogenetically dispersed hosts

The host associations of the fungi isolated from symptomatic seedlings, in combination with the results of our inoculation experiments, provide support for our hypothesis that seedling pathogens in these tropical forests are often multi-host (Figs **1**, **3**; Table **1**). We speculate that our study underestimates the true breadth of the host ranges of the observed multi-host fungi, as we sampled and tested a small fraction of the >824 tree species present locally (Pyke *et al*., 2001). We cannot assess host specialization for the 33 singleton OTUs and some may be strict specialists. Moreover, host-specialized fungi may have gone unobserved due to our culture-based approach, which limited sampling and excluded biotrophic fungi, generally exhibiting greater specificity than necrotrophs (Agrios, 2005). That said, multi-host pathogens of seedlings, leaves, and seeds were also observed in previous Panama-based studies (Augspurger & Wilkinson, 2007; Gilbert & Webb, 2007; Schweizer *et al*., 2013; Sarmiento *et al*., 2017).

In general, a given fungal OTU was isolated from heterofamilial tree species, host use was phylogenetically overdispersed, and pairwise similarities in community composition of seedling-associated fungi were unrelated to pairwise phylogenetic distances between tree species (Figs **1**, **2**; Tables **S4, S5**). Together these results suggest that, contrary to our second hypothesis, there is no phylogenetic signal to host range for these multi-host seedling pathogens. The absence of a significant phylogenetic signal is consistent with one Panama-based study of foliar pathogens (Schweizer *et al*., 2013), but conflicts with other studies in temperate and tropical plant communities (Gilbert & Webb, 2007; Gilbert *et al*., 2012; Parker *et al*., 2015; Chen *et al*., 2019). Our study did not include as many congeneric or confamilial host pairs as Gilbert and Webb (2007) or Gilbert *et al*. (2012), potentially hindering our ability to detect a signal. Alternatively, pathogen host use may track phylogenetically labile plant defenses (Kursar *et al*., 2009) rather than phylogenetically conserved defenses (Fine *et al*., 2006). Parallel evolution of similar defenses in unrelated lineages would explain why the pathogens in this and other studies can infect many distantly related tree species.

The positive relationships between OTU isolation frequency and the number of hosts observed during our survey and inoculation experiments (Fig. **S5**) suggest that host generalists are abundant members of the pathogen community and that, if sampling was expanded, a given fungal OTU would be detected in additional tree species. Many seedling pathogens, including the soil-borne fungi and oomycetes causing seedling damping-off, are passively dispersed (e.g., by splashing rain), with limited dispersal distances (Madden, 1997). Thus, the ability to infect a wide range of hosts, particularly spatially aggregated hosts, would increase a pathogen’s likelihood of persistence. For pathogens with limited dispersal, host use may be more tightly linked with plant habitat associations than phylogenetic relatedness, making them more likely to infect tree species that commonly co-occur spatially and temporally (Roy, 2001). Furthermore, just as selection should favor multi-host pathogens in species-rich assemblages, selection should favor true generalists, whose host ranges are decoupled from plant relatedness, in plant communities that are phylogenetically overdispersed on the small spatial scale relevant to dispersal-limited pathogens (Swenson *et al*., 2007; Swenson *et al*., 2012; Pearse *et al*., 2013).

### Tree species share pathogens, but are differentially impacted

A given plant species is often infected by multiple microorganisms and these microorganisms are frequently shared within a plant community (Hersh *et al*., 2012; Sarmiento *et al*., 2017; Spear & Mordecai, 2018). We observed overlap among the fungal communities of distantly related tree species (Fig. **1**; Table **S5**), a pattern likely to intensify with increased sampling. The presence of a particular fungus in the seedlings of multiple tree species does not mean that those hosts are equally likely to be infected by that fungus. Pathogens may disproportionally infect certain hosts based on availability, defense traits, and/or habitat (Ferrer & Gilbert, 2003).

Disease progression among susceptible hosts is mediated by multiple factors, including defense traits. Based on our inoculation experiments, tree species were differentially impacted by shared pathogens (Fig. **3**; Table **1**), in support of our third hypothesis and consistent with the host-specific impacts of multi-host pathogens previously observed (Davidson *et al*., 2000; Augspurger & Wilkinson, 2007; Hersh *et al*., 2012; Sarmiento *et al*., 2017). Most of the tree species we tested were seemingly resistant to the 10 pathogenic isolates (Fig. **3**), but that does not preclude their susceptibility to numerous untested fungi. Moreover, these tree species may actually be susceptible to infection by the pathogenic isolates in our study, but their resistance mechanisms prevent extensive pathogen colonization and symptom development. In which case, these resistant tree species may act as reservoir hosts that facilitate the persistence of pathogens within the community (Haas *et al*., 2011).

### Plant traits and disease susceptibility

Disease pressure varies spatially, particularly across environmental gradients (Augspurger, 1984a; Brenes-Arguedas *et al*., 2009; Defossez *et al*., 2011; Haak *et al*., 2012; Hersh *et al*., 2012; Spear *et al*., 2015). For plants adapted to areas with low and/or inconsistent disease pressure, such as areas with higher light, lower humidity, and inconsistent precipitation (Augspurger, 1984a; Spear *et al*., 2015), the allocation costs of physical and chemical defenses may outweigh the benefits and vice versa for plants adapted to areas with high and/or consistent disease pressure (Coley & Barone, 1996; Strauss & Agrawal, 1999; Endara & Coley, 2010). Consistent with variable selective pressures, and previous field and greenhouse studies (Augspurger, 1984b; Augspurger & Kelly, 1984; McCarthy-Neumann & Kobe, 2008), seedlings of shade-intolerant tree species and tree species restricted to, or more abundant in, drier forests (dry-site species) were more likely to suffer disease than seedlings of shade-tolerant and wet-site tree species in our inoculation experiments (Fig. **4**; Tables **S6, S7**). Engelbrecht *et al*. (2007) found no correlation between a species’ light requirements and its occurrence in drier versus wetter sites, but evergreen trees are more common in wetter sites, while deciduous trees dominate drier sites (Santiago *et al*., 2004). Shade-tolerant and evergreen tree species, both with long-lived leaves, tend to invest more in pest defenses than shade-intolerant and deciduous species (Coley & Barone, 1996; Santiago *et al*., 2004).

For shade-tolerant, shade-intolerant, and dry-site tree species, the probability of disease decreases with increasing seed size (Fig. **4**). This is perhaps unsurprising as large-seeded species can invest more in defenses and better compensate for lost tissue (Armstrong & Westoby, 1993; Green & Juniper, 2004). Our results are inconsistent with two Panama-based studies (Augspurger, 1984a; Augspurger & Kelly, 1984), which found no correlation between seed size and pathogen susceptibility, possibly because they only included wind-dispersed species while our analyses included wind- and vertebrate-dispersed species (average seed dry masses: 1.9 - 686 versus 3 - 2334 mg).

### Consequences for plant communities

Neither disturbance nor microhabitat specialization are considered sufficient to maintain the hyperdiversity of tropical forests (Leigh *et al*., 2004). While host-specialized pests are commonly touted as the drivers of diversity maintenance (Leigh *et al*., 2004; Comita *et al*., 2014), we show that generalist pathogens are frequently responsible for seedling death and disease. The host-specific impacts of multi-host pathogens and differences among tree species in pathogen susceptibility observed by this and other studies (Augspurger & Wilkinson, 2007; Hersh *et al*., 2012; Spear *et al*., 2015) suggest that generalized pathogens may also influence plant community composition by unevenly affecting seedling recruitment. First, variability in host competence may maintain plant community diversity by creating a dilution effect (Ostfeld & Keesing, 2000). Overall disease risk should be lowered if the most competent host(s) is/are rare within the community because pathogen transmission would be slowed between individuals of susceptible species. Accordingly, the presence of less competent hosts should protect poorly defended host species from pathogen-caused local extinction. Second, generalist pathogens with host-specific impacts may alter direct and indirect competition among co-occurring species (Alexander & Holt, 1998; Barrett *et al*., 2009; Mordecai, 2011). Host-generalized pathogens could promote coexistence if a competition-defense tradeoff exists among plant species and superior competitors are more susceptible (Mordecai, 2011).

Finally, interspecific variation in host susceptibility to generalist pathogens may maintain diversity because disease-sensitive trees (e.g., shade-intolerant and dry-site spp.) are excluded from disease-prone habitats (e.g., wetter, aseasonal forests and low-light understories). By reinforcing resource partitioning and limiting the geographic distributions of species, generalist pathogens contribute to the spatial turnover of plant species and the maintenance of local and regional forest diversity.

## Acknowledgments

This research was funded by Simons Foundation Award 429440, Center for Tropical Forest Science, Smithsonian Tropical Research Institute, Garden Club of America, University of Utah’s Global Change and Sustainability Center, and A. Herbert and Marian W. Gold Scholarship. Additional logistical support and facilities were provided by the Smithsonian Tropical Research Institute, S. Mangan, and the Arnold Lab at University of Arizona. We thank J. Sertich, D. Martinez, B. Wolfe, J. Ibarra, H. Espinosa, C. Swatek, M. Schmoker, D. Sandberg, J. Shaffer, and C. Gonzalez for field and lab assistance; R. Perez for help with tree species identification; J. Wright, G. Gilbert, S. Mangan, A. E. Arnold, S. Higginbotham, T. Kursar, and P. Coley for advice; T. Brenes-Arguedas for sharing her culture collection, and N. Anaya and G. LaRosa, who contributed to that collection; Parque Natural Metropolitano, D. Roubik, Barro Colorado Nature Monument, and the Ministerio de Ambiente for authorizing this research; and four anonymous reviewers for comments that improved this manuscript.

## Author Contributions

ERS designed and conducted the research, analyzed the data, and wrote the first draft of the manuscript, ERS and KDB contributed to manuscript revisions and approved the final version.

## Supporting Information

**Fig. S1** Examples of disease in the forests of Panama.

**Fig. S2** Details of shadehouse-based inoculation experiments.

**Fig. S3** Rank abundance plot and OTU accumulation curve.

**Fig. S4** Overlap in fungal OTUs among sampling years, methods used to obtain symptomatic seedlings, isolation media, and tissue sampled.

**Fig. S5** Correlation between OTU host range and isolation frequency.

**Table S1** Taxonomic assignments, traits, sampling effort, and observed OTUs for tree species evaluated in our survey and experiments.

**Table S2** Methodological details pertaining to the multi-year collection of symptomatic seedlings, and microbial isolation and sequencing.

**Table S3** Average light levels, air temperatures, and relative humidities of the shadehouses used for the inoculation experiments versus ambient conditions.

**Table S4** Estimated taxonomic placement, isolation frequency, number of observed hosts, estimated host specialization, and phylogenetic pattern of host use of the OTUs.

**Table S5** Overlap in seedling-associated OTUs among tree species.

**Table S6** Results of the beta-binomial generalized linear regression with the proportion of diseased seedlings as a function of seed size and shade tolerance.

**Table S7** Average estimates based on the best-ranked beta-binomial generalized linear regressions with the proportion of diseased seedlings as a function of seed size and spatial distribution relative to annual rainfall.

**Methods S1** Methods used to estimate the taxonomic placements of the 66 OTUs and assign nomenclature.

