## Supporting Information for "Host-generalist fungal pathogens of seedlings may maintain forest diversity via host-specific impacts and differential susceptibility among tree species"

The following Supporting Information is available for this article:

**Fig. S1** Examples of disease symptoms in the forests of Panama.

**Fig. S2** Details of shadehouse-based inoculation experiments.

**Fig. S3** Rank abundance plot and OTU accumulation curve.

**Fig. S4** Overlap in fungal OTUs among sampling years, methods used to obtain symptomatic seedlings, isolation media, and tissue sampled.

**Fig. S5** Correlation between OTU host range and isolation frequency.

**Table S1** Taxonomic assignments, traits, sampling effort, and observed OTUs for tree species evaluated in our survey and experiments.

**Table S2** Methodological details pertaining to the multi-year collection of symptomatic seedlings, and microbial isolation and sequencing**.**

**Table S3** Average light levels, air temperatures, and relative humidities of the shadehouses used for inoculation experiments versus ambient conditions.

**Table S4** Estimated taxonomic placement, isolation frequency, number of observed hosts, estimated host specialization, and phylogenetic pattern of host use of the OTUs.

**Table S5** Overlap in seedling-associated OTUs among tree species**.**

**Table S6** Results of the beta-binomial generalized linear regression with the proportion of diseased seedlings as a function of seed size and shade tolerance.

**Table S7** Average estimates based on the best-ranked beta-binomial generalized linear regressions with the proportion of diseased seedlings as a function of seed size and spatial distribution relative to annual rainfall.

**Methods S1** Methods used to estimate the taxonomic placement of the 66 OTUs and assign nomenclature.

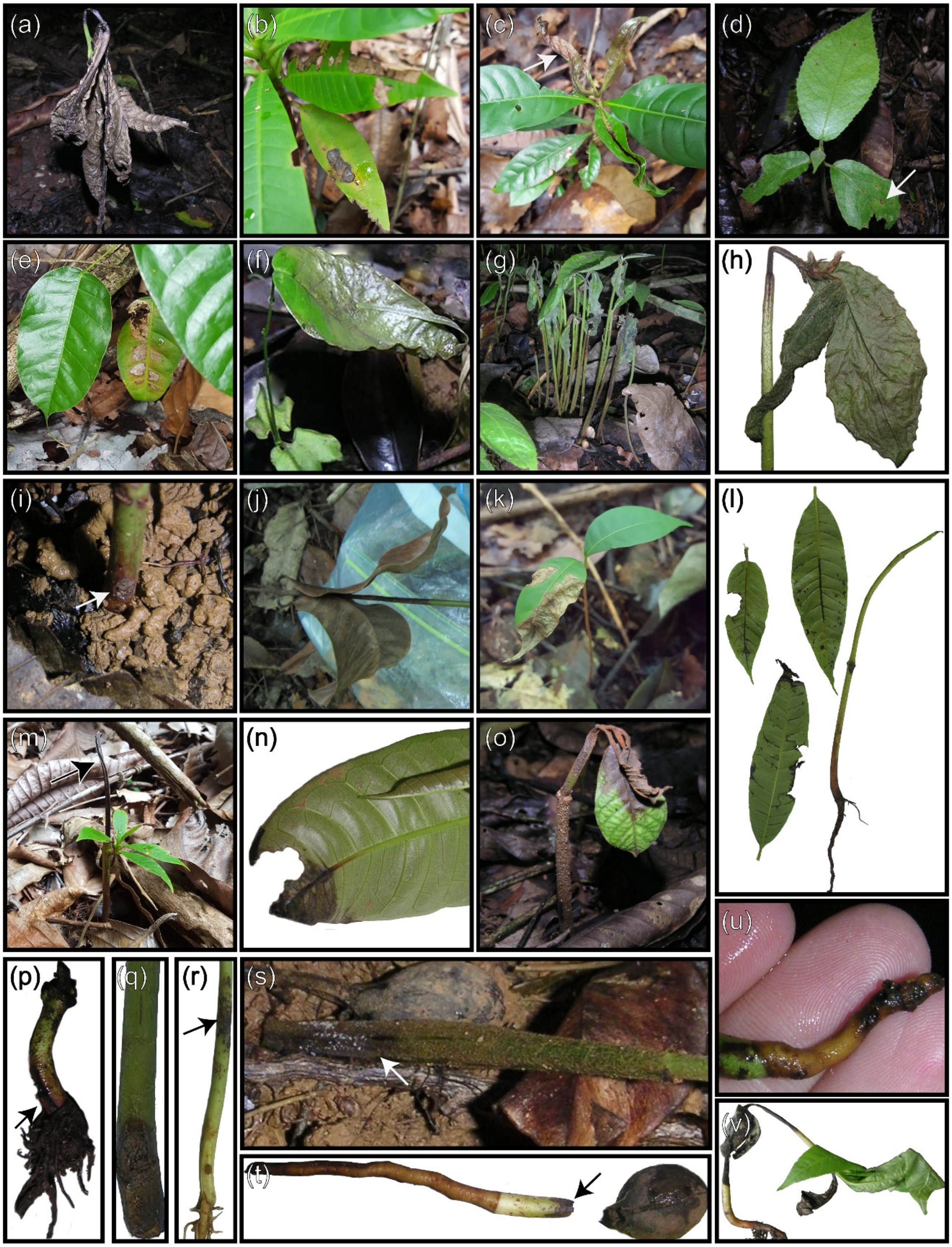


**Fig. S1** Disease symptoms on the (a,g-j,m,o,p-t,v) stems, (b-f,k,l,n,o) leaves, and (u) root of seedlings in the forests of Panama. In some panels, arrows direct the viewer's attention to disease.

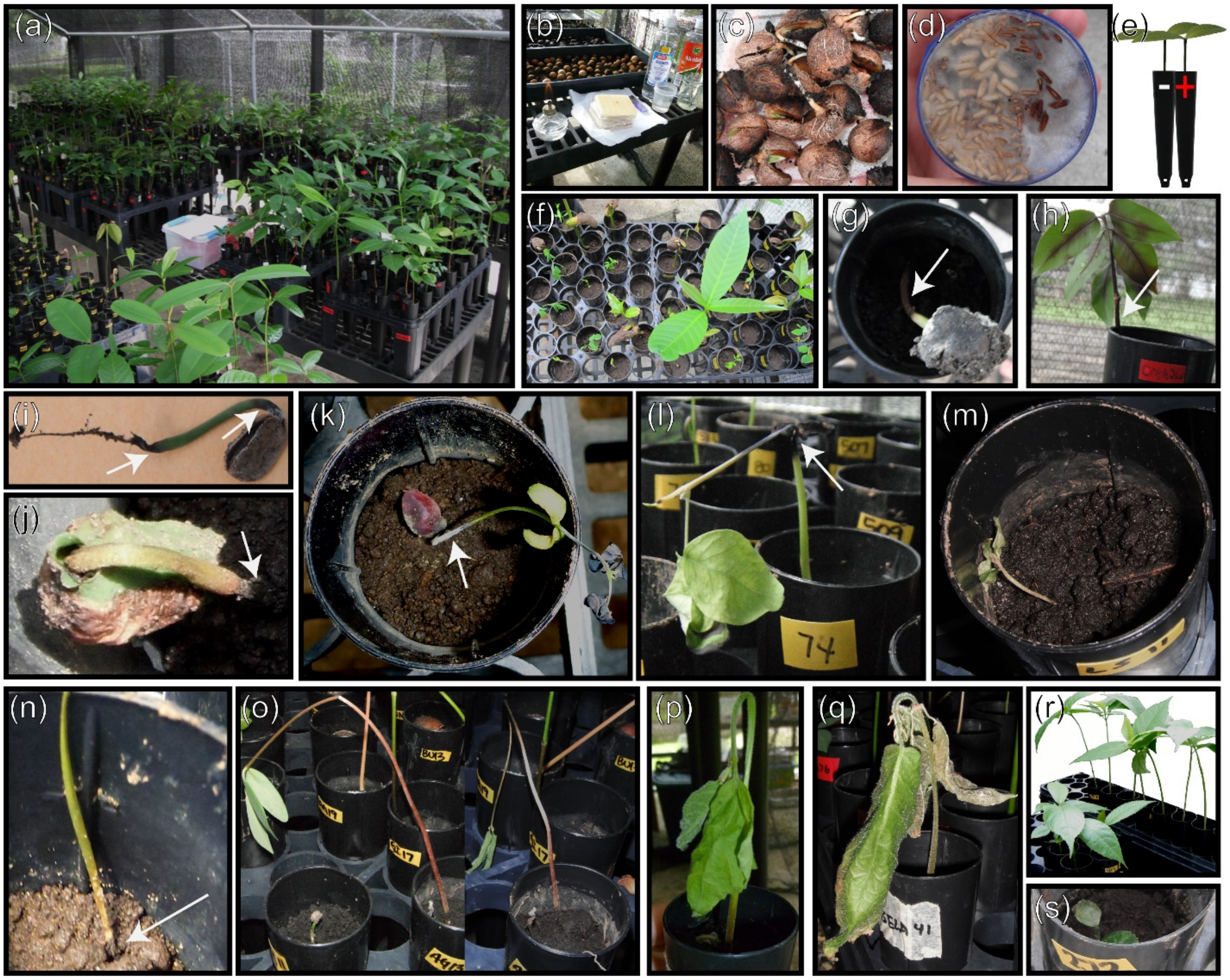


**Fig. S2** (a) Inoculation experiments were conducted in Smithsonian Tropical Research Institute shadehouses in Gamboa, Panama. (b) Surface-sterilized seeds were germinated in flats of autoclave-sterilized commercial soil. (c,f) Seedlings were transplanted to individual pots containing autoclaved commercial soil and (d,e) either rice visibly colonized by one of the fungal isolates or inoculum-free, autoclave-sterilized rice. Disease was documented every 3 d and was categorized as mortality, (g-o) stem damage, (p,q) wilting, and (r,s) stunting.

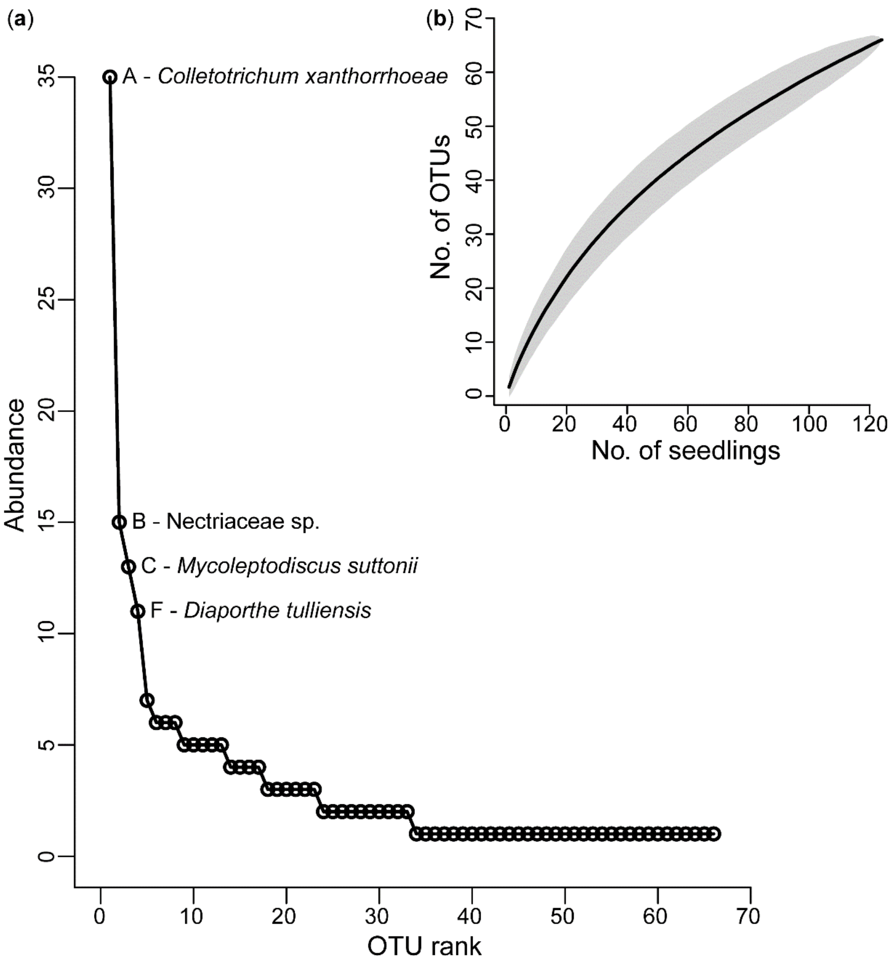


**Fig. S3** (a) The 66 observed OTUs are ranked from most to least abundant on the horizontal axis, with the total number of isolates per OTU plotted on the vertical axis (full dataset) (BiodiversityR package; Kindt & Coe, 2005). Most of the OTUs are rare (50% singletons), indicated by the steep shape of the curve. The four common OTUs (observed >10 times and comprising 35% of the isolates) are named. (b) Non-asymptotic accumulation of OTUs isolated from 124 symptomatic seedlings (full dataset) (vegan package; Oksanen et al., 2019). The curve, derived from the observed richness and representing the mean accumulation of OTUs over 999 randomizations of seedling order, indicates incomplete sampling and a diverse community.

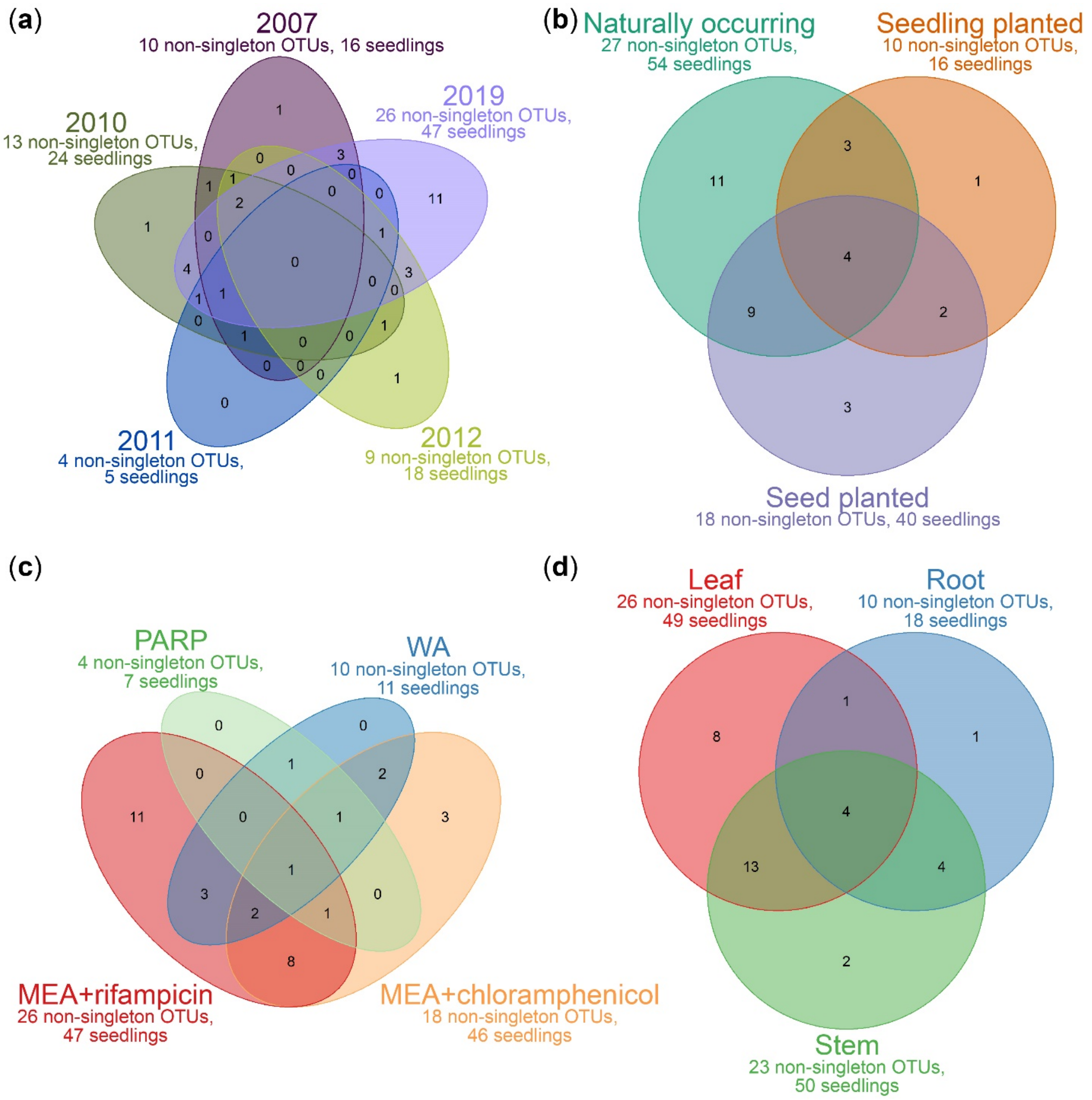


**Fig. S4** Venn diagrams depicting the overlap in non-singleton fungal operational taxonomic units (OTUs) among the (a) five sampling years, (b) three methods used to obtain seedlings with disease, (c) four media used for isolation, and (d) three tissues sampled (data subset A) (VennDiagram package; Chen, 2018). (a) Fungi, and two oomycetes, were isolated from symptomatic seedlings in Panama over five years. (b) Symptomatic seedlings were obtained in three ways: (i) opportunistic collection of naturally occurring seedlings, (ii) seedlings germinated in a shadehouse and then transplanted to forest sites, and (iii) surface-sterilized seeds planted directly in forest sites. (c, d) The advancing margin(s) of diseased area(s)

was/were excised, and the excised tissue piece(s) (leaf, stem, and/or root) was/were surface sterilized (Gilbert & Webb, 2007) and plated on (i) Water Agar (WA), (ii) Pimaricin, Ampicillin, Rifampicin, and Pentachloronitrobenzene (PARP); and/or Malt Extract Agar (MEA) amended with antibiotic to prevent bacterial growth, either (iii) chloramphenicol or (iv) rifampicin. See Table S2 and Spear (2007) for additional methodological details. (a) Of the 33 non-singleton OTUs, 19 were observed in more than one year. While no OTUs were observed across all five years, three OTUs were observed across four of the sampling years. The greatest number of unique, non-singleton OTUs was observed in 2019, the year we collected the greatest number of seedlings. (b) Eighteen non-singleton OTUs were isolated from seedlings obtained using more than one method. Four non-singleton OTUs were isolated from seedlings obtained using all three methods. We isolated the greatest number of unique, non-singleton OTUs from naturally occurring seedlings, the most common sampling method. (c) Nineteen non-singleton OTUs were isolated on multiple media. One non-singleton OTU was isolated from tissue pieces plated on all four media. We isolated the greatest number of unique, non-singleton OTUs on MEA+rifampicin, the medium used for the greatest number of seedlings and tissue pieces. (d) Twenty-two non-singleton OTUs were isolated from multiple tissues. Four non-singleton OTUs were isolated from all three tissues. We isolated the greatest number of unique, non-singleton OTUs from leaves, the best-sampled tissue.

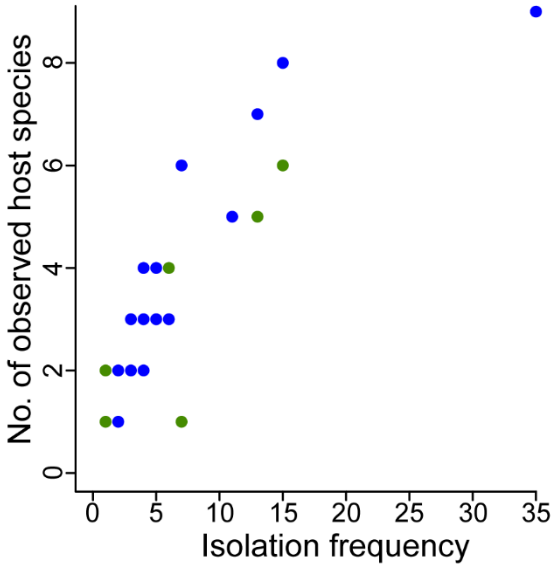


**Fig. S5** The observed host range of an OTU is positively correlated with isolation frequency (survey-based assessment of host range: blue points, one-tailed Spearman's rank correlation rho = 0.96, $P < 0.001$; host range observed during the inoculation experiments: green points, one-tailed Spearman's rank correlation rho = 0.72, $P = 0.053$), suggesting that multi-host fungi may be common in this system.

**Table S1** Tree species from which putative pathogens were isolated (original host = OH, 26 tree spp.) and/or for which vulnerability to pathogens was assessed (target = T, 35 tree spp.) via inoculation experiments. For each tree species, the following is listed: a two- or three-letter code (for Tables 1, S2, and S5), taxonomic assignments, and the number of seedlings collected, sites from which seedlings were collected, unique isolates observed, and OTUs observed. Average seed dry mass (mg), shade tolerance, and spatial distribution relative to annual rainfall are listed for the tree species used to explore the relationship between disease susceptibility and plant life history traits. The imperfect match between original hosts and targets was driven by space, time, and seed availability constraints. Additionally, several tree species that were not original hosts were included in the inoculation experiments as phytometers (measures of isolate pathogenicity) because of previously observed disease susceptibility (e.g., L. seemannii).

| **Role** | **Species** | **Code** | **Family** | **Order** | **Seed mass (mg)**[1] | **Shade tol.**[2] | **Dist.**[3] | **Seedlings collected** | **Sites** | **Isolates** | **OTUs** |
|---|---|---|---|---|---|---|---|---|---|---|---|
| OH, T | *Anacardium excelsum* (Bertero ex Kunth) Skeels | ANE | Anacardiaceae | Sapindales | 1507 | | dry | 20 | 5 | 41 | 24 |
| OH, T | *Dalbergia retusa* Hemsl. | DR | Fabaceae | Fabales | 130 | tol | dry | 18 | 2 | 20 | 10 |
| OH | *Pouteria reticulata* (Engl.) Eyma | PR | Sapotaceae | Ericales | | | | 11 | 1 | 30 | 18 |
| OH, T | *Virola surinamensis* (Rol. ex Rottb.) Warb | VS | Myristicaceae | Magnoliales | | | | 11 | 3 | 15 | 11 |
| OH | *Faramea occidentalis* (L.) A.Rich. | FO | Rubiaceae | Gentianales | | | | 7 | 1 | 13 | 7 |
| OH | *Protium panamense* (Rose) I.M.Johnst. | PP | Burseraceae | Sapindales | | | | 7 | 1 | 18 | 11 |
| OH | *Protium tenuifolium* (Engl.) Engl. | PT | Burseraceae | Sapindales | | | | 6 | 2 | 10 | 9 |
| OH, T | *Calophyllum longifolium* Willd. | CL | Calophyllaceae | Malpighiales | | | | 5 | 1 | 14 | 9 |
| OH, T | *Castilla elastica* Cerv. | CE | Moraceae | Rosales | 203.4 | | dry | 5 | 2 | 6 | 4 |
| OH, T | *Hymenaea courbaril* L. | HC | Fabaceae | Fabales | | | | 5 | 3 | 6 | 5 |
| OH | *Cassia moschata* Kunth | CAM | Fabaceae | Fabales | | | | 4 | 3 | 5 | 4 |
| OH, T | *Lacmellea panamensis* (Woodson) Markgr. | LAP | Apocynaceae | Gentianales | 237.4 | tol | wet | 4 | 3 | 7 | 5 |
| OH | *Nectandra cuspidata* Nees & Mart. | NC | Lauraceae | Laurales | | | | 4 | 2 | 5 | 5 |
| OH, T | *Cochlospermum vitifolium* (Willd.) Spreng. | CV | Bixaceae | Malvales | 26 | intol | | 3 | 1 | 4 | 4 |
| OH | *Swietenia macrophylla* King | SWM | Meliaceae | Sapindales | | | | 2 | 1 | 2 | 2 |
| OH, T | *Trichilia tuberculata* (Triana & Planch.) C. DC. | TT | Meliaceae | Sapindales | 151 | tol | | 2 | 2 | 2 | 2 |
| OH, T | *Brosimum utile* (Kunth) Oken | BU | Moraceae | Rosales | 1763.5 | | wet | 1 | 1 | 2 | 2 |
| OH, T | *Cojoba rufescens* (Benth.) Britton & Rose | CR | Fabaceae | Fabales | 236.2 | tol | dry | 1 | 1 | 2 | 2 |
| OH | *Dipteryx oleifera* Benth. | DO | Fabaceae | Fabales | | | | 1 | 1 | 1 | 1 |

| | | | | | | | | | | | |
|---|---|---|---|---|---|---|---|---|---|---|---|
| OH, T | *Genipa americana* L. | GA | Rubiaceae | Gentianales | 123 | tol | dry | 1 | 1 | 1 | 1 |
| OH | *Mouriri myrtilloides* (Sw.) Poir. | MM | Melastomataceae | Myrtales | | | | 1 | 1 | 1 | 1 |
| OH | *Ormosia coccinea* (Aubl.) Jacks. | OC | Fabaceae | Fabales | | | | 1 | 1 | 1 | 1 |
| OH, T | *Ormosia macrocalyx* Ducke | OM | Fabaceae | Fabales | 379 | tol | dry | 1 | 1 | 1 | 1 |
| OH | *Randia armata* (Sw.) DC. | RA | Rubiaceae | Gentianales | | | | 1 | 1 | 1 | 1 |
| OH | *Spondias mombin* L. | SPM | Anacardiaceae | Sapindales | | | | 1 | 1 | 2 | 2 |
| OH, T | *Tetragastris panamensis* (Engl.) Kuntze | TEP | Burseraceae | Sapindales | 295 | tol | | 1 | 1 | 1 | 1 |
| T | *Annona glabra* L. | AG | Annonaceae | Magnoliales | 229 | intol | wet | | | | |
| T | *Coccoloba manzinellensis* Beurl. | COM | Polygonaceae | Caryophyllales | | | | | | | |
| T | *Copaifera aromatica* Dwyer | CA | Fabaceae | Fabales | 893.6 | tol | dry | | | | |
| T | *Eugenia nesiotica* Standl. | EN | Myrtaceae | Myrtales | 346.5 | tol | | | | | |
| T | *Garcinia intermedia* (Pittier) Hammel | GI | Clusiaceae | Malpighiales | 541 | tol | | | | | |
| T | *Guapira standleyana* Woodson | GS | Nyctaginaceae | Caryophyllales | 61.2 | tol | | | | | |
| T | *Inga goldmanii* Pittier | IG | Fabaceae | Fabales | | | | | | | |
| T | *Inga sapindoides* Willd. | IS | Fabaceae | Fabales | 406.4 | intol | | | | | |
| T | *Jacaranda copaia* (Aubl.) D.Don | JC | Bignoniaceae | Lamiales | 5 | intol | | | | | |
| T | *Lacistema aggregatum* (P.J.Bergius) Rusby | LA | Lacistemataceae | Malpighiales | 10.7 | intol | | | | | |
| T | *Licania platypus* (Hemsl.) Fritsch | LIP | Chrysobalanaceae | Malpighiales | | | | | | | |
| T | *Luehea seemannii* Triana & Planch | LS | Malvaceae | Malvales | 3 | intol | dry | | | | |
| T | *Pachira quinata* (Jacq.) W.S.Alverson | PQ | Malvaceae | Malvales | 40 | intol | dry | | | | |
| T | *Posoqueria latifolia* (Rudge) Schult. | POL | Rubiaceae | Gentianales | 196.5 | tol | | | | | |
| T | *Psychotria limonensis* K.Krause | PSL | Rubiaceae | Gentianales | 6.6 | tol | | | | | |
| T | *Psychotria marginata* Sw. | PM | Rubiaceae | Gentianales | 7.5 | tol | | | | | |
| T | *Quararibea asterolepis* Pittier | QA | Malvaceae | Malvales | 335 | tol | | | | | |
| T | *Siparuna pauciflora* (Beurl.) A. DC. | SP | Siparunaceae | Laurales | | | | | | | |
| T | *Swartzia simplex* (Sw.) Spreng. | SS | Fabaceae | Fabales | 1025 | tol | | | | | |
| T | *Symphonia globulifera* L.f. | SG | Clusiaceae | Malpighiales | 2334 | tol | wet | | | | |
| T | *Tocoyena pittieri* (Standl.) Standl. | TOP | Rubiaceae | Gentianales | 956.2 | tol | wet | | | | |

[1]Seed mass sources: (1) **Daws MI, Garwood NC, Pritchard HW. 2005.** Traits of recalcitrant seeds in a semi-deciduous tropical forest in Panama: some ecological implications. *Functional Ecology* **19**: 874–885. (2) **Myers JA, Kitajima K. 2007.** Carbohydrate storage enhances seedling shade and stress tolerance in a neotropical forest. *Journal of Ecology* **95**: 383–395. (3) ER Spear, unpublished data. (4) **Svenning JC, Wright SJ. 2005.** Seed limitation in a Panamanian forest. *Journal of Ecology* **93**: 853–862. (5) **Wright SJ, Kitajima K, Kraft NJB, Reich PB, Wright IJ, Bunker DE, Condit R, Dalling JW, Davies SJ, Díaz S *et al.* 2010.** Functional traits and the growth–mortality trade-off in tropical trees. *Ecology* **91**: 3664–3674.

[2]Shade tolerance sources: (1) **Augspurger CK. 1984.** Light requirements of neotropical tree seedlings: a comparative study of growth and survival. *Journal of Ecology* **72**: 777–795. (2) **Brown SH, Mark S. 2013.** Fact sheet: *Annona glabra*. USDA, Cooperative Extension Service, University of Florida, IFAS, Florida A. & M.

[WWW document] URL http://www.doc-developpement-durable.org/file/Arbres-Fruitiers/FICHES_ARBRES/Cachiman-cochon-mammier-Annona-glabra/Pond%20Apple%20-%20Lee%20County%20Extension%20-%20University%20of%20Florida.pdf. [accessed 21 May 2020]. (3) **Comita LS, Aguilar S, Pérez R, Lao S, Hubbell SP. 2007.** Patterns of woody plant species abundance and diversity in the seedling layer of a tropical forest. *Journal of Vegetation Science* **18**: 163–174. (4) **Hall JS, Ashton MS. 2016.** *Guide to early growth and survival in plantations of 64 tree species native to Panama and the Neotropics*. Balboa, Ancón, República de Panamá: Smithsonian Tropical Research Institute. (5) **Kitajima K, Llorens AM, Stefanescu C, Timchenko MV, Lucas PW, Wright SJ. 2012.** How cellulose-based leaf toughness and lamina density contribute to long leaf lifespans of shade-tolerant species. *New Phytologist* **195**: 640–652. (6) **Martin WA, Flores EM. 2002.** *Copaifera aromatica* Dwyer. In Vozzo JA, ed. *Tropical tree seed manual.* Washington DC, USA: USDA Forest Service, 405–407. (7) **Molofsky J, Augspurger CK. 1992.** The effect of leaf litter on early seedling establishment in a tropical forest. *Ecology* **73**: 68–77. (8) **Paul GS, Montagnini F, Berlyn G P, Craven DJ, van Breugel M, Hall JS. 2012.** Foliar herbivory and leaf traits of five native tree species in a young plantation of Central Panama. *New Forests* **43**: 69–87. (9) **Pearcy RW, Valladares F, Wright SJ, De Paulis EL. 2004.** A functional analysis of the crown architecture of tropical forest *Psychotria* species: do species vary in light capture efficiency and consequently in carbon gain and growth? *Oecologia* **139**: 163–177.

[3]Distribution sources: (1) **Engelbrecht BM, Comita LS, Condit R, Kursar TA, Tyree MT, Turner BL, Hubbell SP. 2007.** Drought sensitivity shapes species distribution patterns in tropical forests. *Nature* **447**: 80–82. (2) **Condit R, Pérez R, Daguerre N. 2010.** *Trees of Panama and Costa Rica*. Princeton, NJ, USA: University Press. (3) **Perez R, Condit R. 2020.** Tree Atlas of Panama. [WWW document] URL http://ctfs.si.edu/webatlas/maintreeatlas.php. [accessed 21 May 2020]

**Table S2** Methodological details pertaining to the multi-year collection of symptomatic seedlings, and microbial isolation and sequencing. Seedlings were collected from the lowland tropical forests of Panama in 2007, 2010-2012, and 2019 by E. R. Spear and T. Brenes-Arguedas (column 1). Symptomatic seedlings were obtained by opportunistically collecting naturally occurring seedlings and by baiting pathogens from the soil by planting seedlings or surface-sterilized seeds directly in the forest sites (col. 2). The forest sites from which seedlings were collected included: Buena Vista Peninsula (BV) and Barro Colorado Island (BCI) in Barro Colorado Nature Monument, Gunn Hill in Ciudad del Saber (formerly Fort Clayton; FC), private property on Santa Rita Ridge (SRR), Parque Natural Metropolitano (PNM), and Sendero Camino de Cruces (CC) and Sendero del Charco in Parque Nacional Soberanía (SC) (col. 3; see Fig. S1 in Spear (2017) for a map and additional site details). In total, 124 seedlings of 26 tree species were collected, but the tree species collected varied by site and year (col. 4; see Table S1 for full species names). Seedlings were collected during the rainy season (col. 5 provides date ranges). For all 124 collected seedlings, the advancing margin(s) of diseased area(s) was/were excised, and the excised symptomatic tissue piece(s) was/were surface sterilized following Gilbert & Webb (2007) prior to plating. Fungi (and two oomycetes) were isolated on four media: Water Agar (WA; 20 isolates); Pimaricin, Ampicillin, Rifampicin, and Pentachloronitrobenzene (PARP†; 11 isolates); and Malt Extract Agar (MEA) amended with antibiotic to prevent bacterial growth, either chloramphenicol (chloram; 59 isolates) or rifampicin (rifamp; 121 isolates) (col. 6). Morphologically unique isolates were subcultured into pure culture, and a piece of mycelium was excised from each pure culture for molecular analysis (col. 6). The DNA extractions, PCR amplifications, and bidirectional Sanger sequencing‡ of either the nuclear ribosomal internal transcribed spacer (ITS) region (149 isolates) or the ITS plus an adjacent portion of the large subunit (ITS+LSU) (62 isolates) were completed over multiple years and by multiple labs: the lab of A. E. Arnold at the University of Arizona (methods followed lab's protocols; e.g., Sandberg et al., 2014); the International Cooperative Biodiversity Groups (ICBG) lab at the Smithsonian Tropical Research (STRI) (methods followed lab's protocols; e.g., Higginbotham et al., 2014); the molecular research lab at STRI Naos Marine Laboratories; the lab of K. D. Broders at STRI; and Macrogen, Inc. (col. 7). The primers ITS1F, ITS5, ITS4, and LR3 (Vilgalys & Hester, 1990; White et al., 1990; Gardes & Bruns, 1993) were used for amplification and sequencing (col. 8). Edited DNA sequences for all, but one, of the fungal isolates collected from 2007-2012 have been deposited in NCBI GenBank (col. 9). Edited DNA sequences for one fungal isolate collected in 2011, and 119 fungal isolates and two oomycetes collected in 2019 will be deposited upon manuscript acceptance (col. 9). All 211 isolates (from 2007-2019) were used for our survey-based assessment of the host associations and ranges of putative pathogens (col. 10). Twenty-seven of the isolates collected in 2010 and 2011 were used for our experimental assessments of pathogenicity and host range conducted in 2011 and 2012 (col. 11).

| **Year & collector** | **Source of symptomatic seedlings** | **Forest site(s)** | **No. of seedlings & tree spp. collected** | **Date range of seedling collection** | **No. of unique isolates, media used, plant tissue sampled** | **Facilities where extractions, amplifications, & sequencing were completed** | **Primers used** | **NCBI GenBank Accession numbers** | **Isolates used for survey-based assessment of the host associations & ranges of putative pathogens** | **Isolates used for experimental assessments of pathogenicity & host range** |
|---|---|---|---|---|---|---|---|---|---|---|
| 2007§<br>Brenes-Arguedas | Seeds were germinated in a STRI shadehouse and then seedlings were planted in forest sites | (1) BV<br>(2) FC<br>(3) SRR | 22<br><br>Tree spp:<br>(1) BU,<br>(2) CAM,<br>(3) CV,<br>(4) HC,<br>(5) LAP,<br>(6) NC,<br>(7) OC,<br>(8) OM<br>(9) PT,<br>(10) SWM | Jul. 21-Nov.13, 2007 | 28 fungal isolates<br><br>Media:<br>(1) WA,<br>(2) PARP†<br><br>Plant tissue:<br>(1) stem,<br>(2) root | (1) Arnold Lab | ITS5, ITS1F, ITS4, LR3 | KY413686-KY413775 | Yes | No |
| 2010§<br>Spear | (1) Surface-sterilized seeds planted directly in forest sites (see Spear et al. 2015 for additional details)<br>(2) Naturally occurring seedings | (1) SRR<br>(2) PNM | 28<br><br>Tree spp:<br>(1) ANE,<br>(2) CE,<br>(3) CR,<br>(4) GA,<br>(5) HC,<br>(6) PT,<br>(7) RA,<br>(8) TEP,<br>(9) TT,<br>(10) VS | Jul 20-Nov. 16, 2010 | 34 fungal isolates<br><br>Media:<br>(1) MEA+chloram<br>(2) PARP†<br><br>Plant tissue:<br>(1) stem,<br>(2) root,<br>(3) leaf | (1) Arnold Lab,<br>(2) STRI ICBG lab,<br>(3) STRI mol. research lab | ITS1F, ITS4, LR3 | | Yes | Yes |

| | | | | | | | | | | |
|---|---|---|---|---|---|---|---|---|---|---|
| 2011§ Spear | Naturally occurring seedings | (1) PNM<br>(2) BCI<br>(3) CC<br>(4) SC | 8<br><br>Tree spp:<br>(1) ANE,<br>(2) CL,<br>(3) DO | May 21-Jun. 9, 2011 | 8 fungal isolates<br><br>Media:<br>(1) MEA+chloram<br>Plant tissue:<br>(1) stem,<br>(2) root,<br>(3) leaf | (1) STRI ICBG lab,<br>(2) STRI mol. research lab,<br>(3) Broders Lab,<br>(4) Macrogen, Inc. | ITS5, ITS1F, ITS4, LR3 | | Yes | Yes |
| 2012§ Spear | Surface-sterilized seeds were planted directly in forest sites | (1) SRR<br>(2) PNM | 18<br><br>Tree sp:<br>(1) DR | Jul. 5-19, 2012 | 20 fungal isolates<br><br>Media:<br>(1) MEA+chloram<br><br>Plant tissue:<br>(1) stem,<br>(2) root,<br>(3) leaf | (1) Arnold Lab | ITS5, LR3 | | Yes | No |
| 2019 Spear | Naturally occurring seedings | (1) BCI | 48<br><br>Tree spp:<br>(1) ANE,<br>(2) CL,<br>(3) FO,<br>(4) LP,<br>(5) MM,<br>(6) PP,<br>(7) PR,<br>(8) PT,<br>(9) SPM,<br>(10) VS | Sept. 27-Oct. 14, 2019 | 119 fungal isolates & 2 oomycetes<br><br>Media:<br>(1) MEA+rifamp<br><br>Plant tissue:<br>(1) stem,<br>(2) root,<br>(3) leaf | (1) STRI mol. research lab,<br>(2) Broders Lab¶,<br>(3) Macrogen, Inc. | ITS5, ITS4 | Will be deposited upon manuscript acceptance | Yes | No |

† While PARP contains antifungals and was used with the intention of isolating oomycetes, only fungi were cultivated. We believe there was a problem with the antifungals or medium preparation. All non-singleton fungal operational taxonomic units isolated on PARP were also isolated on MEA+rifamp, MEA+chloram, and/or WA (Fig. **S4c**).

‡ For 17 isolates, paired-end reads were not possible due to a low quality read in one direction.

§ Methods and sequences previously published in Spear (2007).

¶ Amplified DNA was generated by either direct colony polymerase chain reaction (DC-PCR; following Walch *et al.*, 2016) or PCR of DNA extracted in TE (Tris-EDTA) buffer. A T100™ Thermal Cycler (Bio-Rad Laboratories, Inc, Hercules, CA, USA) was used for amplification: 3 min of initial denaturation at 95 °C, followed by 36 cycles of 95°C for 30 s, 54°C for 30 s, and 72 °C for 1 min, and a final extension step of 72°C for 10 min (modified from U'Ren *et al.*, 2010). Amplification was verified with gel electrophoresis and GelRed® Nucleic Acid Gel Stain (Biotium, Inc., Fremont, CA, USA).

**Table S3** Average light levels, air temperatures, and relative humidities of the two shadehouses used for the inoculation experiments (Fig. S2) versus the forest understory. The screened-in shadehouses are located in Gamboa, Panama (9°7'9.87"N, 79°42'4.96"W). The photosynthetically active radiation (PAR, in µmol of photons s-1 m-2) reaching shadehouse seedlings was measured during the afternoon of a uniformly overcast day (Oct. 12, 2011). Measurements were taken inside and directly outside the shadehouses with a LI-250 light meter, a LI-190 quantum sensor, and a one-meter LI-191 line quantum sensor (LI-COR, Lincoln, NE, USA). The mean (± SD) wet season forest understory light value was obtained from Brenes-Arguedas et al. (2011). In 2006 and 2007, Brenes-Arguedas et al. (2011) measured instantaneous light 0.5 m above the forest floor with LI-190 quantum sensors (LI-COR), and QSO sensors (Apogee Instruments, Logan, UT, USA) and CR200 and CR1000 data-loggers (Campbell Scientific, Inc., Logan, UT, USA) in a nearby (within c. 16 km) forest in Barro Colorado Nature Monument (BCNM), Buena Vista Peninsula (9°11'N, 79°49'W). In each shadehouse, air temperature and relative humidity (RH) were measured at 10-min intervals (CS500 probe, Campbell Scientific). Hourly mean temperature and minimum and maximum RH were recorded on a CR200 datalogger (Campbell Scientific). Measurements were taken November 3-5 and 5-11, 2011 for shadehouses 2 and 1, respectively. We obtained forest understory air temperature and RH data for the same time period (Nov. 3-11, 2011) from the Physical Monitoring Program of the Smithsonian Tropical Research Institute (Paton, 2019a,b) Forest understory air temperature and RH data were measured at 15-min intervals with a CS215 Temperature and Relative Humidity Probe (Campbell Scientific) at a nearby (within c. 16 km) forest in BCNM, the Lutz Tower (sensor height 1m) on Barro Colorado Island (9°9'42.06"N, 79°50'15.83"W). We calculated hourly means from STRI's timestamped 15-min interval data to allow for comparison with our air temperature and RH data. Both shadehouses were used for the inoculation experiments in 2011 and only shadehouse 1 was used in 2012.

| Location | Mean ± SD<br>% of full PAR | Mean ± SD air temperature‡ | Mean min. ± SD RH‡ | Mean max. ± SD RH‡ |
|---|---|---|---|---|
| Shadehouse 1 | 1.4% † | 26.1 ± 1.9ºC$^{ns}$ | 85.1 ± 8.6%*** | 88.1 ± 6.4%*** |

| | | | | |
|---|---|---|---|---|
| Forest understory | 1.3 ± 0.8% | 25.4 ± 0.9ºC | 95.6 ± 0.5% | 98.3 ± 1.2% |
| Shadehouse 2 | 1.7% † | 25.6 ± 1.3ºC$^{ns}$ | 87.6 ± 6.9%*** | 90.5 ± 4.6%*** |
| Forest understory | 1.3 ± 0.8% | 25.3 ± 0.5ºC | 96.7 ± 0.6% | 99.0 ± 0.4% |

†Because inside and outside PAR measurements were not taken simultaneously, we calculated mean % of full PAR as the average of the PAR values recorded inside (shadehouse 1: *n* = 13, shadehouse 2: *n* = 8) divided by the average of the PAR values recorded outside (shadehouse 1: *n* = 9, shadehouse 2: *n* = 8) and we could not calculate SD. For this reason, and because we did not have access to the raw data summarized in Brenes-Arguedas *et al.* (2011), we did not use statistical tests to compare shadehouse and forest understory light levels.

‡ The air temperature and relative humidity data are not normally distributed (assessed by variable and shadehouse with Shapiro-Wilk tests of normality [stats package; R Core Team, 2020], all $P < 0.01$). Therefore, two-tailed Wilcoxon rank-sum tests (stats package; R Core Team, 2020) were used to compare the air temperature and RH conditions of the two shadehouses to those of the forest understory (ns, not significant; ***, $P < 0.001$; Quinn & Keough, 2002). *n* = 150 per group for shadehouse 1 and its corresponding understory data. *n* = 46 per group for shadehouse 2 and its corresponding understory data.

**R Core Team. 2020.** R: a language and environment for statistical computing. R Foundation for Statistical Computing, Vienna, Austria.

**Table S4** For each of the 66 operational taxonomic units (OTUs), we report its estimated taxonomic placement and the UNITE database accession code(s) associated with the reference sequence(s) used to assign nomenclature (Kõljalg et al., 2013), the number of times it was isolated, and the number of tree species from which it was isolated. For the 33 non-singleton OTUs (data subset A), we report estimated host specialization based on the d' index (value and category; low (L): 0–0.33, moderate (M): 0.34–0.67) (bipartite package; Dormann et al., 2008). To determine if there is a phylogenetic signal to the host range of the 31 OTUs isolated from multiple tree species, we used the ses.mpd function (picante package; Kembel et al., 2010). In that analysis, the observed mean phylogenetic distance (MPD) between tree species infected by a given OTU is compared to the MPD expected under a null model with random host-OTU associations (full dataset; 999 permutations). Negative and positive standardized effect sizes indicate phylogenetic clustering and phylogenetic overdispersion of host use, respectively (i.e., a given OTU associates with hosts more closely [Obs. MPD < Exp. MPD] or more distantly [Obs. MPD > Exp. MPD] related than expected by chance) (Kembel et al., 2010). P-values <0.05 and >0.95 indicate significant phylogenetic clustering and overdispersion, respectively (Kembel, 2010). Significant and marginally significant P-values are in bold text and marked with a letter indicating the phylogenetic pattern of host use (clustering, C; overdispersion, O).

| **OTU–Est. taxonomic placement - UNITE code(s)†** | **No. of isolates** | **No. of obs. host spp.** | **Est. host specialization: d' index & category** | **Obs. MPD** | **Null model mean MPD** | **Null model SD of MPD** | **Standardized effect size** | ***P*** |
|---|---|---|---|---|---|---|---|---|
| A–*Colletotrichum xanthorrhoeae* - SH1543705 (*C. xanthorrhoeae*), SH1543739 (unidentified) | 35 | 9 | 0.225 - L | 219.3 | 233.0 | 21.2 | -0.645 | 0.34 |
| B–Nectriaceae sp. - SH1610517 (Nectriaceae), SH1610162 (*Cylindrocladium buxicola*) | 15 | 8 | 0.292 - L | 278.8 | 233.0 | 23.3 | 1.969 | **0.97**[O] |
| C–*Mycoleptodiscus suttonii* - SH1562616 | 13 | 7 | 0.538 - M | 262.4 | 233.8 | 26.8 | 1.068 | 0.85 |
| F–*Diaporthe tulliensis* - SH1540611 | 11 | 5 | 0.347 - M | 223.3 | 234.7 | 34.7 | -0.328 | 0.49 |
| I–Sordariomycetes sp. - SH1541118 | 7 | 6 | 0.402 - M | 230.6 | 235.0 | 29.2 | -0.149 | 0.56 |
| G–*Clonostachys rosea* - SH1522825 | 6 | 4 | 0.274 - L | 280.0 | 232.4 | 39.4 | 1.209 | 0.81 |
| D–*Cylindrocladiella variabilis* - SH1610166 | 6 | 3 | 0.350 - M | 307.8 | 230.5 | 48.6 | 1.591 | 0.85 |

| | | | | | | | | |
|---|---|---|---|---|---|---|---|---|
| H–Xylariaceae sp. - SH1541124 | 6 | 3 | 0.242 - L | 234.4 | 235.0 | 49.2 | -0.013 | 0.60 |
| AAD–*Mycoleptodiscus suttonii* - SH1562616 | 5 | 4 | 0.272 - L | 233.9 | 232.5 | 40.0 | 0.035 | 0.66 |
| AU–*Beltrania pseudorhombica* - SH1563660 | 5 | 4 | 0.061 - L | 299.2 | 232.2 | 38.4 | 1.744 | **0.95** [O] |
| J–*Pseudopestalotiopsis theae* - SH1552673 | 5 | 4 | 0.206 - L | 279.8 | 233.4 | 39.9 | 1.162 | 0.78 |
| L–*Neopestalotiopsis foedans* - SH1552672 (*N. foedans*), SH2700365 (Pezizomycotina) | 5 | 4 | 0.031 - L | 217.5 | 234.1 | 40.0 | -0.416 | 0.41 |
| M–*Lasiodiplodia gonubiensis* - SH1507365 | 5 | 3 | 0.126 - L | 320.3 | 232.1 | 47.3 | 1.866 | **0.94** [O] |
| S–*Colletotrichum citricola* - SH1543707 | 4 | 4 | 0.198 - L | 233.9 | 233.1 | 38.6 | 0.021 | 0.66 |
| N–*Gliocladiopsis elghollii* - SH1546330 | 4 | 3 | 0.669 - M | 204.0 | 235.6 | 51.0 | -0.619 | 0.30 |
| E–*Beltrania pseudorhombica* - SH1563660 | 4 | 2 | 0.350 - M | 238.3 | 232.2 | 68.4 | 0.090 | 0.56 |
| O–*Fusarium pseudensiforme* - SH1546322 (*F. pseudensiforme*), SH1546498 (Hypocreales) | 4 | 2 | 0.593 - M | 235.1 | 233.9 | 67.4 | 0.017 | 0.36 |
| AV–*Cylindrocladiella variabilis* - SH1212036 | 3 | 3 | 0.122 - L | 320.3 | 232.9 | 48.1 | 1.817 | **0.97** [O] |
| AW–*Calonectria pseudonaviculata* - SH1610162 | 3 | 3 | 0.118 - L | 237.2 | 232.4 | 47.3 | 0.103 | 0.64 |
| AX–Nectriaceae sp. - SH1212162 (*Neonectria* sp.), SH1610423 (*Cylindrocladiella* sp.), SH1212036 (*C. variabilis*) | 3 | 3 | 0.511 - M | 237.2 | 234.6 | 49.4 | 0.053 | 0.67 |
| AY–*Beltraniella* cf. *endiandrae* - SH1563659 | 3 | 3 | 0.383 - M | 196.6 | 233.7 | 50.1 | -0.741 | 0.17 |
| P–*Colletotrichum thailandicum* - SH1543711 | 3 | 3 | 0.083 - L | 237.2 | 234.8 | 50.2 | 0.048 | 0.76 |
| Q–*Diaporthe* cf. *columnaris* - SH1540633 (Diaporthales), SH1540609 (*D. columnaris*) | 3 | 2 | 0.226 - L | 119.7 | 232.1 | 68.4 | -1.644 | **0.09** [C] |
| AAA–*Diaporthe fraxini-angustifoliae* - SH1540607 | 2 | 2 | 0.481 - M | 235.1 | 235.3 | 71.9 | -0.003 | 0.49 |
| AAB–*Trichoderma spirale* - SH1567965 (*T. spirale*), SH1552633 (Pezizomycotina) | 2 | 2 | 0.156 - L | 215.2 | 232.8 | 67.1 | -0.262 | 0.24 |
| AAC–Xylariaceae sp. - SH1541166 | 2 | 2 | 0.380 - M | 361.3 | 234.9 | 68.6 | 1.841 | 0.87 |
| AZ–*Clonostachys* cf. *miodochialis* - SH1522826 (Hypocreales), SH1522827 (*C. miodochialis*) | 2 | 2 | 0.530 - M | 214.0 | 233.4 | 69.5 | -0.280 | 0.18 |
| K–*Trichoderma spirale* - SH1567965 | 2 | 2 | 0.570 - M | 238.3 | 234.1 | 71.1 | 0.059 | 0.84 |
| R–*Diaporthe fraxini-angustifoliae* - SH1540607 | 2 | 2 | 0.086 - L | 119.7 | 232.1 | 68.4 | -1.644 | **0.09** [C] |
| U–*Colletotrichum magnisporum* - SH1543718 | 2 | 2 | 0.118 - L | 238.3 | 232.6 | 70.1 | 0.082 | 0.67 |
| V–*Xylaria multiplex* - SH1541132 | 2 | 2 | 0.134 - L | 238.3 | 233.6 | 65.6 | 0.073 | 0.73 |
| ‡T–*Macrophomina* - SH1507375 (Botryosphaeriaceae), SH1507369 (*M. phaseolina*) | 2 | 1 | 0.343 - M | | | | | |
| ‡X–*Ceratobasidium* sp. - SH1551758 | 2 | 1 | 0.164 - L | | | | | |
| AA–*Diaporthe endophytica* - SH1540603 | 1 | 1 | | | | | | |

| | | |
|---|---|---|
| AAE–*Colletotrichum thailandicum* - SH1543711 | 1 | 1 |
| AAF–Nectriaceae sp. - SH1610517 | 1 | 1 |
| AAG–*Colletotrichum brevisporum* - SH1543708 | 1 | 1 |
| AAH–*Beltraniella endiandrae* - SH1563659 | 1 | 1 |
| AAI–*Diaporthe* cf. *columnaris* - SH1540609 | 1 | 1 |
| AAJ–*Gliocladiopsis elghollii* - SH1546330 | 1 | 1 |
| AAK–*Cylindrocladiella variabilis* - SH1212036 | 1 | 1 |
| AAL–Sordariomycetes sp. - SH1198320 | 1 | 1 |
| AAM–*Talaromyces* - SH1516144 | 1 | 1 |
| AAN–*Cylindrocladiella variabilis* - SH1212036 | 1 | 1 |
| AB–*Diaporthe siamensis* - SH1540610 | 1 | 1 |
| AC–*Nigrospora* cf. *oryzae* - SH1549605 (unidentified fungus), SH1549606 (*N. oryzae*) | 1 | 1 |
| AD–*Gliocladiopsis* sp. - SH1546383 | 1 | 1 |
| AE–*Pestalotiopsis* sp. - SH1563667 | 1 | 1 |
| AF–*Fusarium equiseti* - SH1610158 | 1 | 1 |
| AG–*Pestalotiopsis rhododendri* - SH1563658 | 1 | 1 |
| AH–Oomycota sp. | 1 | 1 |
| AI–*Gliocephalotrichum cylindrosporum* - SH1546353 | 1 | 1 |
| AJ–*Beltraniopsis neolitseae* - SH1563663 | 1 | 1 |
| AK–*Diaporthe endophytica* - SH1540603 | 1 | 1 |
| AL–*Digitiseta multidigitata* - SH1546391 | 1 | 1 |
| AM–*Endomelanconiopsis endophytica* - SH1507376 | 1 | 1 |
| AN–*Phytophthora palmivora* | 1 | 1 |
| AO–*Xylaria* sp. - SH1554119 | 1 | 1 |
| AP–*Diaporthe endophytica* - SH1540603 | 1 | 1 |
| AQ–*Ramularia* sp. - SH1209934 | 1 | 1 |
| AR–Mycosphaerellaceae sp. - SH1606644 | 1 | 1 |
| AS–*Fusarium* - SH1610159 | 1 | 1 |
| AT–Chaetosphaeriaceae sp. - SH1168551 | 1 | 1 |
| W–Sordariomycetes sp. - SH1198320 | 1 | 1 |
| Y–*Talaromyces marneffei* - SH1516144 | 1 | 1 |
| Z–*Colletotrichum brevisporum* - SH1543708 | 1 | 1 |

†All UNITE Species Hypothesis accession codes, beginning with acronym SH, end in '.08FU' (not shown), denoting the version number and the acronym for Fungi (Kõljalg *et al*., 2013). Because our OTU grouping strategy (99% sequence similarity) often differed from that of the UNITE database (97–100% similarity), multiple, distinct OTUs in our study sometimes share the same nomenclature based on the species hypothesis (SH) of the best-matching reference sequence in UNITE (e.g., OTUs D, AV, AAK, and AAN).

‡There are no ses.mpd results for T - *Macrophomina* sp. and X - *Ceratobasidium* sp. because those OTUs were only isolated from a single host species.

**Table S5** Overlap in operational taxonomic units (OTUs) among tree species, considering only the 13 tree species and 22 OTUs with 3 or more observations (data subset B). Gray cells contain the number of observed OTUs and, in parentheses, isolates collected for each tree species (original hosts in Table S1). White cells contain the shared richness of OTUs between pairs of tree species and, in parentheses, the estimated OTU community similarity (1 – Chao index, which is abundance-based and adjusted for unseen species), ranging from zero (no similarity) to one (identical communities). Black cells contain the pooled number of OTUs for each pair of tree species.

| | ANE | CL | CM | CE | CV | DR | FO | HC | LP | PR | PP | PT | VS |
|---|---|---|---|---|---|---|---|---|---|---|---|---|---|
| ANE | 13 (29) | 3 (0.30) | 0 (0) | 1 (0.03) | 2 (0.35) | 4 (0.47) | 3 (0.34) | 0 (0) | 2 (0.36) | 7 (0.81) | 4 (0.36) | 2 (0.23) | 7 (1) |
| CL | 17 | 7 (12) | 0 (0) | 0 (0) | 1 (0.10) | 2 (0.24) | 2 (0.35) | 0 (0) | 2 (0.33) | 7 (1) | 3 (0.42) | 2 (0.32) | 2 (0.17) |
| CM | 15 | 9 | 2 (3) | 1 (0.24) | 0 (0) | 2 (0.54) | 0 (0) | 1 (0.24) | 0 (0) | 1 (0.04) | 0 (0) | 0 (0) | 2 (0.18) |
| CE | 14 | 9 | 3 | 2 (4) | 0 (0) | 1 (0.11) | 0 (0) | 1 (0.21) | 0 (0) | 0 (0) | 0 (0) | 0 (0) | 1 (0.09) |
| CV | 15 | 10 | 6 | 6 | 4 (4) | 1 (0.08) | 0 (0) | 1 (0.21) | 3 (1) | 1 (0.08) | 0 (0) | 0 (0) | 1 (0.16) |
| DR | 16 | 12 | 7 | 8 | 10 | 7 (16) | 2 (0.32) | 1 (0.11) | 1 (0.07) | 4 (0.63) | 2 (0.23) | 1 (0.18) | 2 (0.15) |
| FO | 14 | 9 | 6 | 6 | 8 | 9 | 4 (10) | 0 (0) | 0 (0) | 2 (0.32) | 1 (0.27) | 2 (0.61) | 1 (0.07) |
| HC | 16 | 10 | 4 | 4 | 6 | 9 | 7 | 3 (4) | 1 (0.16) | 0 (0) | 0 (0) | 1 (0.18) | 1 (0.09) |
| LP | 15 | 9 | 6 | 6 | 5 | 10 | 8 | 6 | 4 (6) | 2 (0.21) | 1 (0.13) | 0 (0) | 1 (0.08) |
| PR | 19 | 13 | 14 | 15 | 16 | 16 | 15 | 16 | 15 | 13 (25) | 5 (0.59) | 3 (0.42) | 6 (1) |
| PP | 15 | 10 | 8 | 8 | 10 | 11 | 9 | 9 | 9 | 14 | 6 (13) | 1 (0.21) | 1 (0.06) |
| PT | 15 | 9 | 6 | 6 | 8 | 10 | 6 | 6 | 8 | 14 | 9 | 4 (5) | 1 (0.14) |
| VS | 15 | 14 | 9 | 10 | 12 | 14 | 12 | 11 | 12 | 16 | 14 | 12 | 9 (13) |

**Table S6** Results of the beta-binomial (logit link) generalized linear regression with the proportion of seedlings with disease as a function of seed size and shade tolerance (26 tree species, 179 observations: 42 for intolerant spp., 137 for tolerant spp.). Estimates and standard errors (SE) are log odds of disease for a one-unit increase in the variable. The intercept represents shade-intolerant tree species at a hypothetical seed dry mass (mg) of zero. Model averaging was not done because no other model had a ΔAICc ⩽ 2. P-values indicating statistical significance are in bold (P < 0.05).

| Variable | Estimate | SE | z value | *P* |
|---|---|---|---|---|
| intercept | -0.812 | 0.342 | -2.377 | **0.018** |
| seed size | -0.004 | 0.001 | -3.057 | **0.002** |
| shade tolerance | -1.131 | 0.385 | -2.938 | **0.003** |
| seed size:shade tolerance | 0.004 | 0.001 | -2.457 | **0.014** |

**Table S7** Average estimates based on the best-ranked (ΔAICc ⩽ 2) beta-binomial (logit link) generalized linear regressions with the proportion of seedlings with disease as a function of seed size and spatial distribution relative to annual rainfall (15 tree species, 154 observations: 108 for dry-site spp., 46 for wet-site spp.). Estimates and standard errors are log odds of disease for a one-unit increase in the variable. The intercept represents a dry-site tree species at a hypothetical seed dry mass (mg) of zero. P-values indicating statistical significance are in bold (P < 0.05).

| Variable | Estimate | Std. error | z value | *P* |
|---|---|---|---|---|
| intercept | -1.517 | 0.222 | 6.829 | **<0.001** |
| seed size | -0.0005 | 0.0003 | 1.593 | 0.111 |
| distribution | -1.255 | 0.619 | 2.028 | **0.042** |
| seed size:distribution | 0.0008 | 0.0006 | 1.286 | 0.198 |

**Methods S1** Methods used to estimate the taxonomic placement of the 66 operational taxonomic units (OTUs) and assign nomenclature.

We estimated the taxonomic placement of the fungi isolated from symptomatic seedlings by querying all 209 fungal sequences in our study against the well-curated and annotated Full UNITE+INSD dataset for Fungi (v. 8.2, released 2020-02-04; Abarenkov *et al*., 2020). Fungal sequences accessioned in the UNITE database have been clustered into what are hypothesized to be species-level groups (Species Hypotheses, SHs), each with a unique accession code. We implemented BLASTN (Altschul *et al*., 1990) in Python with default parameters (blastn -query Spear.fasta -task blastn -db uniteDB -out Python_blastout_1Oct20.txt -evalue 10 -outfmt '6 qseqid pident qcovs evalue bitscore length sallseqid' -max_target_seqs 5 -num_threads 16). For each query, we considered the top five hits and an OTU was assigned nomenclature based on the reference sequence meeting the following criteria: an alignment length ≥75 bp (range 374–1148 bp, mean 759 bp, median 618 bp), an E-value ≤ $10^{-36}$ (all <$10^{-150}$) (as in Radujković *et al*., 2019), query cover ≥90% (range 91–100%, mean 99.5%, median 100%), and the highest percent identity for the OTU (range 93–100%, mean 99.7%, median 100%) (as suggested by Lücking *et al*., 2020).

Following the aforementioned steps of our initial, Python-based approach, there was taxonomic ambiguity for 18 of the 64 fungal OTUs. These OTUs had multiple (2-3) different SH accession codes among the hits with a percent identity of 100% and/or they were not identified to the genus level based on the SH of their 'best' hit. In an attempt to resolve ambiguities and achieve the greatest taxonomic resolution possible, we queried one to two representative sequence(s) from each of those 18 OTUs against the web-based UNITE database (v. 8.2, accessed 9-Oct and 16-Nov-2020; Nilsson *et al.*, 2018), reviewed the top 30 hits for each sequence, and, when possible and appropriate (e.g., no taxonomic conflicts among the named matches), revised their nomenclature based on the criteria specified above (e.g., OTU AC was revised from 'unidentified fungus' to *Nigrospora* cf. *oryzae*). As a result, we manually assigned the nomenclature for 10 of the 18 OTUs with taxonomic ambiguity following our Python-based approach. For those 10 OTUs, we list all relevant SH accession codes (i.e., the SH of the 'best' hit and the SH(s) of the reference sequence(s) used to assign nomenclature) in Table **S4**.

For the two oomycetes (OTUs AH and AN), their sequences were queried against GenBank (accessed 8-Oct-2020; Benson *et al*., 2012) and the curated database *Phytophthora*-ID (v. 2.0, accessed 8-Oct-2020; Grünwald *et al*., 2011).

When annotating OTUs based on reference sequences named to the species level, we considered: (i) 99–100% percent identity a positive match between the query and reference sequence; (ii) 97–98.99% percent identity a close match, the observed differences may fall within the variability of the species (5 OTUs; annotated as cf.); and 90–96.99% percent identity to be a positive match at the genus, but not species, level (3 OTUs: T, AS, AAM).

Dual nomenclature for pleomorphic fungi (i.e., different scientific names for the asexual and sexual forms of single species) has been replaced with a single name for a fungal species (Hawksworth *et al*., 2011). For each OTU, we verified the current accepted name by reviewing Index Fungorum (www.indexfungorum.org), Mycobank (Crous *et al*., 2004), and published literature, and we revised the nomenclature for OTU AF (from *Gibberella intricans* to *Fusarium equiseti*; Xia *et al*., 2019), OTU AQ (from *Mycosphaerella* to *Ramularia*; Wijayawardene *et al.,* 2014), and OTU AW (from *Cylindrocladium buxicola* to *Calonectria pseudonaviculata;* Lombard *et al*., 2010). Additionally, we followed the nomenclature of Hernández-Restrepo *et al*. (2019) for the taxonomic ranks of *Mycoleptodiscus*.

All assigned nomenclatures represent estimates due to several limitations associated with our sequence-based approach (Kang *et al*., 2010; Hofstetter *et al.*, 2019; Lücking *et al.*, 2020). While the nuclear ribosomal internal transcribed spacer (ITS) region is the formal barcode for the molecular identification of fungi (Schoch *et al.*, 2012). It is well established that the ITS region is an unreliable barcode for species discrimination for certain taxa (Lücking *et al.*, 2020); for example, *Calonectria* (Liu *et al.*, 2020), *Diaporthe* (Santos *et al*., 2017), *Colletotrichum* (Marin-Felix *et al.*, 2017), and *Penicillium* (Seifert *et al*., 2007). A multi-locus phylogenetic evaluation is required for accurate species identification (*sensu* Santos *et al*., 2017). Furthermore, precision is limited by the incomplete taxonomic and geographic coverage of existing databases, even well-curated ones like UNITE (Kõljalg *et al*., 2013; Lücking *et al.*, 2020). Additionally, fungal sequences accessioned in the UNITE database have been clustered into SHs based on sequence similarity thresholds ranging from 97–100% (Kõljalg *et al*., 2013; Robbertse *et al.*, 2017; Nilsson *et al*., 2018). While there is no single threshold that appropriately addresses lineage-specific intra- versus interspecific variability for all fungi (Nilsson *et al*., 2008), we used a threshold value of 99% sequence similarity to designate OTUs because: (1) 97–98.5% sequence similarity is too relaxed for species delimitation for some taxonomic groups (Garnica *et al.*, 2016), and (2) we are making statements about host specificity so we adopted a stringent similarity threshold that splits rather than lumps, but that accounts for a small amount of sequencing error. Because our grouping strategy (99% sequence similarity) often differed from that of the UNITE database (97–100% similarity), multiple, distinct OTUs in our study sometimes share the same nomenclature based on the SH of the best-matching reference sequence in UNITE (e.g., OTUs D, AV, AAK, and AAN are all annotated as *Cylindrocladiella variabilis*). It should also be noted that, for certain groups of fungi (e.g., *Trichoderma*), UNITE SHs erroneously include distinct species (Robbertse *et al.*, 2017; Lücking *et al.*, 2020).
